# Structural changes in chromosomes driven by multiple condensin motors during mitosis

**DOI:** 10.1101/2022.06.29.498196

**Authors:** Atreya Dey, Guang Shi, Ryota Takaki, D. Thirumalai

**Affiliations:** The University of Texas at Austin

## Abstract

We created a computational framework that describes the loop extrusion (LE) by multiple condensin I and II motors in order to investigate the changes in chromosome organization during mitosis. The theory accurately reproduces the experimentally measured contact probability profiles for the mitotic chromosomes in HeLa and DT40 cells. The rate of loop extrusion is smaller at the start of mitosis and increases as the cells approach the metaphase. The mean loop size generated by condensin II is about six times larger than the ones created by condensin I. The loops, which overlap with each other, are stapled to a central dynamically changing helical scaffold formed by the motors during the LE process. The structures of the mitotic chromosomes, using a data-driven method that uses the Hi-C contact map as input, are best described as random helix perversion (RHP) in which the handedness changes randomly along the scaffold. The extent of propagation in the RHP structures is less in HeLa cells than in the DT40 chromosomes.

## Introduction

Over the last decade a variety of experimental tools have been developed, which have yielded unprecedented glimpses of the nuclear architecture. These methods fall under the rubric Chromosome Conformation Capture techniques (3C, 4C, 5C, Hi-C, and Micro-C) (Van Steensel and Dekker 2010; Dekker et al. 2002; Lieberman-Aiden et al. 2009; Dixon et al. 2012; Rao et al. 2014; Hsieh et al. 2015), which we collectively refer to as Hi-C. The output from a typical Hi-C experiment is the extraction of the average contact map (CM), a matrix whose elements give the relative probability that two loci, separated by the genomic distance (*s*), are in spatial proximity. Although there are nuances in the organization of the genomes that are not fully captured by the Hi-C experiments, the CMs reveal that two major length scales characterize the structures of mammalian interphase chromosomes. On scales on the order of a few Mbps chromosomes likely undergo microphase separation (Jost et al. 2014; Shi, Liu, et al. 2018; Brackley and Marenduzzo 2020; Belaghzal et al. 2021) into two compartments, which are manifested as checkerboard patterns in the ensemble-averaged CM (Lieberman-Aiden et al. 2009). On a scale of ∼ 500 Kbps, there are fine structures in the CMs, which are referred to as Topologically Associating Domains (TADs) (Dixon et al. 2012).

As the cell transitions from interphase to mitosis, the structures of the chromosome change dramatically. The distinct separation between the euchromatin and heterochromatin that is prominent in interphase chromosomes breaks down, and chromosomes condense, adopting a characteristic rod-like or cylindrical shape (T. Cremer and M. Cremer 2010; Manders, Kimura, and Cook 1999; Haaf and Schmid 1991; Marsden and U. Laemmli 1979; Paulson and U. Laemmli 1977; Yokota et al. 1995; Marko 2008). For a comprehensive overview of mitotic chromosomes, we refer the readers to a recent review (Paulson, Hudson, et al. 2021). In an insightful study, Gibcus et al. 2018 performed Hi-C experiments to monitor the changes in the chicken DT40 cell as the cells transition from the G2 to mitotic phase. By building on earlier studies (Green et al. 2012; Hirano and Mitchison 1994; Ono, Fang, et al. 2004b; Gerlich et al. 2006; Dej, Ahn, and Orr-Weaver 2004; Hagstrom et al. 2002; Hudson et al. 2003; Oliveira, Coelho, and Sunkel 2005; Siddiqui et al. 2006; Strunnikov, Hogan, and Koshland 1995), they showed that the changes in the organization of the chicken DT40 genome are driven by ATPase motors condensin I and condensin II. The major findings of this study that are most relevant for the present work are: (i) In less than 10 minutes after the cells enter the prophase, the signatures of compartment formation as well as the characteristics of the TADs are lost. (ii) In the time window, 10 min≤ *t* ≤ 60 min, the most prominent feature in the contact probability *P*(*s*) as a function of the genomic (*s*) distance between two loci, is a peak whose location shifts to higher *s* values as the cell cycle progresses (see Fig 3A in Gibcus et al. 2018). (ii) By successively depleting condensin II (CAP-H2-mAID in Fig 5A in Gibcus et al. 2018) and condensin I (CAP-H-mAID in Fig 5A in Gibcus et al. 2018), it was argued that the presence of the peak and its movement to higher *s* values is due to the emergence of a helical scaffold, and possible change in the helical turn length in the mitotic chromosome. Moreover, they argued that the formation of the helical structure is largely driven by the condensin II motor with condensin I playing little or no role. Informed by the experimental observations (Earnshaw and U.K. Laemmli 1983; Paulson and U. Laemmli 1977; Pienta and Coffey 1984; Paturej et al. 2016) they also performed polymer simulations in which chromosomes were confined to a cylinder, and constrained to adopt a helical scaffold from which non-overlapping loops emanate. The parameters in the polymer model were adjusted to reproduce the *P*(*s*) versus *s* extracted from the Hi-C map.

That the structures of the mitotic chromosome are likely to be helical was proposed long ago (Marsden and U. Laemmli 1979; Paulson and U. Laemmli 1977; Rattner and Lin 1985; Earnshaw 1988; Woodcock, Frado, and Rattner 1984; Ohnuki 1968; Kuwada 1939; De La Tour and U.K. Laemmli 1988). However, the physical mechanisms that drive helix formation are not evident from the Hi-C (Gibcus et al. 2018) or from the more recent imaging experiments (Chu et al. 2020). In particular, there is no understanding of the roles played by multiple copies of the two motor proteins (condensin I and II) in shaping the chromosome structures during mitosis. This is a difficult problem to solve because it involves thousands of motors (see SI for an estimate of the number of motors). Their structures and ATPase cycles (even for single condensin motors) have not been fully elucidated. A potential clue on how to proceed comes from many single molecule experiments, (Ganji et al. 2018 and Kong et al. 2020), which have not only measured the speed of loop extrusion (LE) but have also shown that a single condensin molecule extrudes loops in a one sided asymmetric manner or by two-sided symmetric fashion. The single molecule studies set the stage for our theoretical investigations of the mitotic chromosome architecture involving multiple motors, and how it changes as the cell cycle progresses.

In order to explain the experimental results, we first generalized our (Takaki et al. 2021) theory for loop extrusion (LE) for a single motor to multiple motors, (details in the Methods section and SI) in order to determine changes in the organization of chromosomes as the cell progresses through different stages of the mitosis cycle. The extruded loops were then used in a polymer model based on the Generalized Rouse Model for Chromosomes (GRMC)(Bryngelson and Thirumalai 1996; Shi and Thirumalai 2019; Rouse Jr 1953).The resulting active GRMC (A-GRMC) model *without any adjustable parameter* shows that asymmetric LE alone can create mitotic chromosome structures observed in both HeLa and DT40 cells(Abramo et al. 2019; Gibcus et al. 2018).

In order to reproduce *P*(*s*) at other time-points (*t* = 7.5, 10, 15 and 30 mins) in the cell cycle of the condensin II depleted chicken DT40 cells, we found that it was necessary to vary the step sizes taken by the motor. The A-GRMC does not quantitatively account for the dependence of *P*(*s*) on *s* for the condensin I depleted DT40 cells, which suggests that the coordinated action of condensin II motors in generating long loops has to differ from condensin I, which generates shorter loops on an average. In order to go beyond the A-GRMC, we calculated the three-dimensional (3D) structures of the chromosomes throughout the cell cycle for DT40 cells using the Hi-C-polymer-physics-structures (HIPPS) method (Shi and Thirumalai 2021). The calculated *P*(*s*) using the 3D structures agree *quantitatively* with the experimental data at all times. By calculating the angular correlation function of the structures, we affirm the emergence of helical conformations as the cells transition to the mitotic phase. The 3D structures suggest that the loops in the mitotic chromosomes are arranged around an overall helical scaffold, which changes from one realization to another. The helical regions randomly alternate in chirality, suggestive of **r**andom **h**elix (or tendril) **p**erversion - **RHP**. Thus, through a combination of the A-GRMC theory and 3D structures calculated using only the Hi-C map, we account nearly quantitatively for many (if not all) aspects of the structural changes in chromosomes during mitosis.

## Results

### The A-GRMC simulations quantitatively reproduces *P*(*s*) for mitotic chromosomes

The theory for loop extrusion by multiple condensin motors is based on the mechanism sketched in Fig. 1. The physical picture is based on the scrunching mechanism (Takaki et al. 2021; Ryu et al. 2020), which assumes that the heads of the motors that are bound to the DNA are relatively immobile whereas the hinge moves towards the stationary heads during the catalytic cycle. Loop extrusion occurs by repeated transition of the hinge towards the motor heads through an allosteric mechanism (Takaki et al. 2021). The parameters needed for the A-GRMC simulations (Table 1) are either taken or inferred from experiments (see details in the SI). Fig. 2(A) shows a snapshot of the loop conformation generated by multiple motors in a 10 Mbps chromosome. The helical-like scaffold is generated by the motor proteins to which the loops are stapled. We find evidence for both Z-loop and nested loop (Fig. 2(B)). The loops extruded by condensin I are on an average smaller in size compared to the ones generated by condensin II (Fig. 2(C)). Because the linear density of condensin I is greater than condensin II, the loops extruded by the former resemble a dense polymer brush-like structure. On the other hand, loops generated by condensin II are fewer in number but longer, which facilitates the formation of long-range contacts between the loci. It is worth pointing right at the outset that the helical scaffold, and hence the structures of the mitotic chromosomes, varies from one realization to another. The A-GRMC simulations also show that the majority of the loops are nested with the fraction of Z-loops being less (Fig. 2(D)). Finally, the simulations using the scrunching mechanism quantitatively accounts for the dependence of *P*(*s*) on *s* for HeLa cells and DT40 cells (Fig. 2(E)). The results in Fig. 2 provide a mechanistic basis for generating the dynamical helical scaffold created by multiple condensin motors in several cell types.

**Table 1:** Parameters in the A-GRMC simulations.

|  | condensin I<br>(100 Mbp) | condensin II<br>(100 Mbp) |
| --- | --- | --- |
| Extrusion rate | 1500 bp/s | 1500 bp/s |
| $\tau_P$ | 7 s | 7 s |
| $\tau_B$ | 120 s | 360 s |
| $n_t$ | 2553 | 343 |
| $n_b$ | 1205 | 210 |
| $n_u = n_t - n_b$ | 1348 | 133 |
| $p_{\text{on}}$ | 0.0075 | 0.0044 |
| $p_{\text{off}}$ | 0.0083 | 0.0028 |

**Figure 1:**
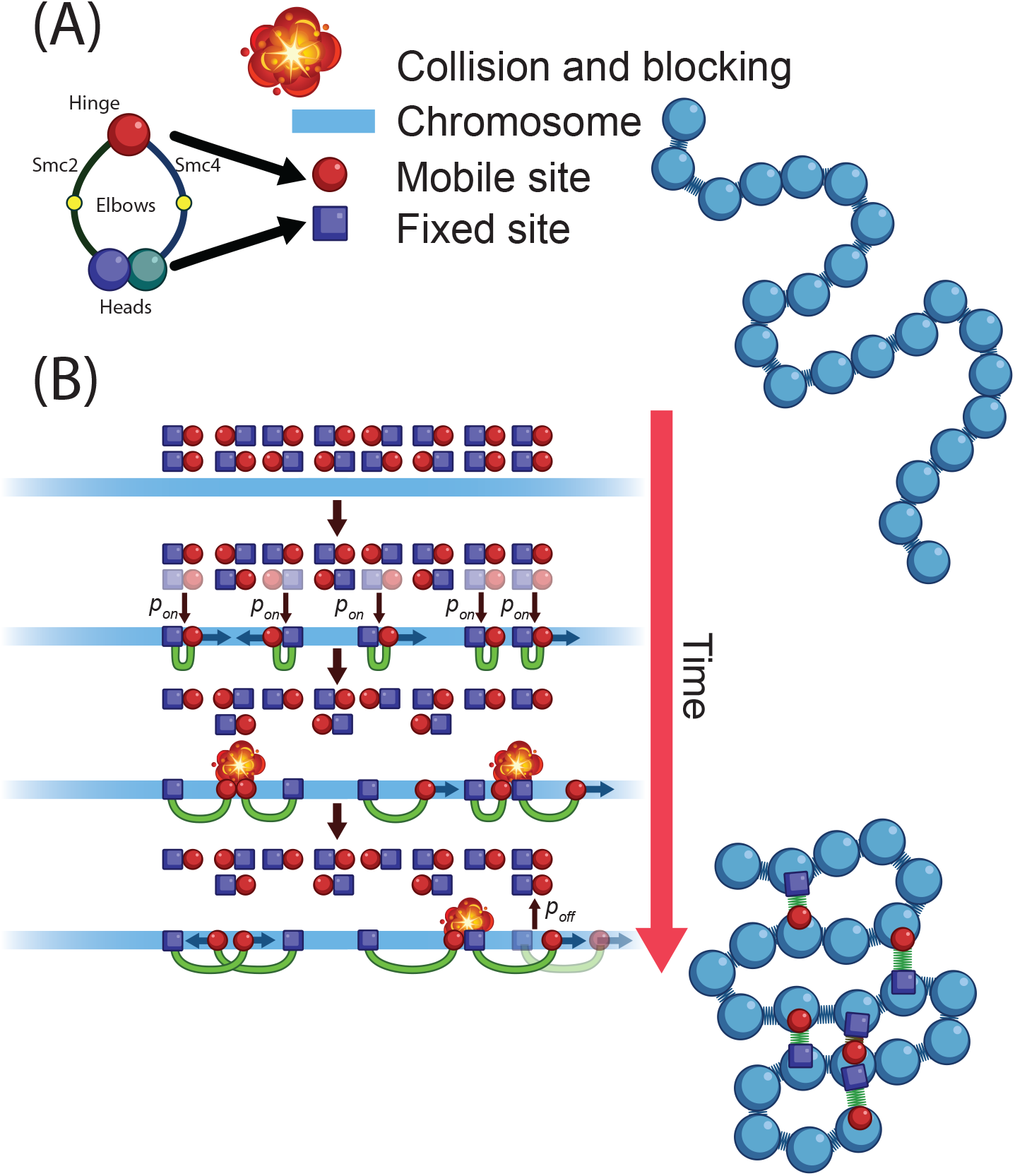
Mechanism of multiple condensin driven loop extrusion using A-GRMC. (A) Schematic illustration of the structural features of condensin (I or II) motor. The representation of the fixed (heads) and mobile (hinge) is based on the scrunching mechanism. The mobile and static sites of condensin are shown as red circles and blue squares, respectively. The dark blue arrows show the direction of loop extrusion by condensin. The green semi-circle represents the extruded loop. Condensin motors bind to DNA with probability *p*_on_ and unbind with probability *p*_off_ shown with black arrows. The mobile site takes a step (shown with small blue arrows) with probability *P*_step_ at each time step. When the motor steps loops are extruded unidirectionally with a step-size that is sampled from Eq. S3 in the SI. Only the mobile site moves along the DNA. The movement is paused for *τ*_*p*_ = 7 seconds if the motor encounters another site (mobile or static). The chromosome, which is divided into 10 Kbps monomers, is shown in blue. Adjacent loci are linked by harmonic springs, and looped loci are connected by harmonic springs as well (green). The mobile and static heads of condensin are shown as red circle and square, respectively.

**Figure 2:**
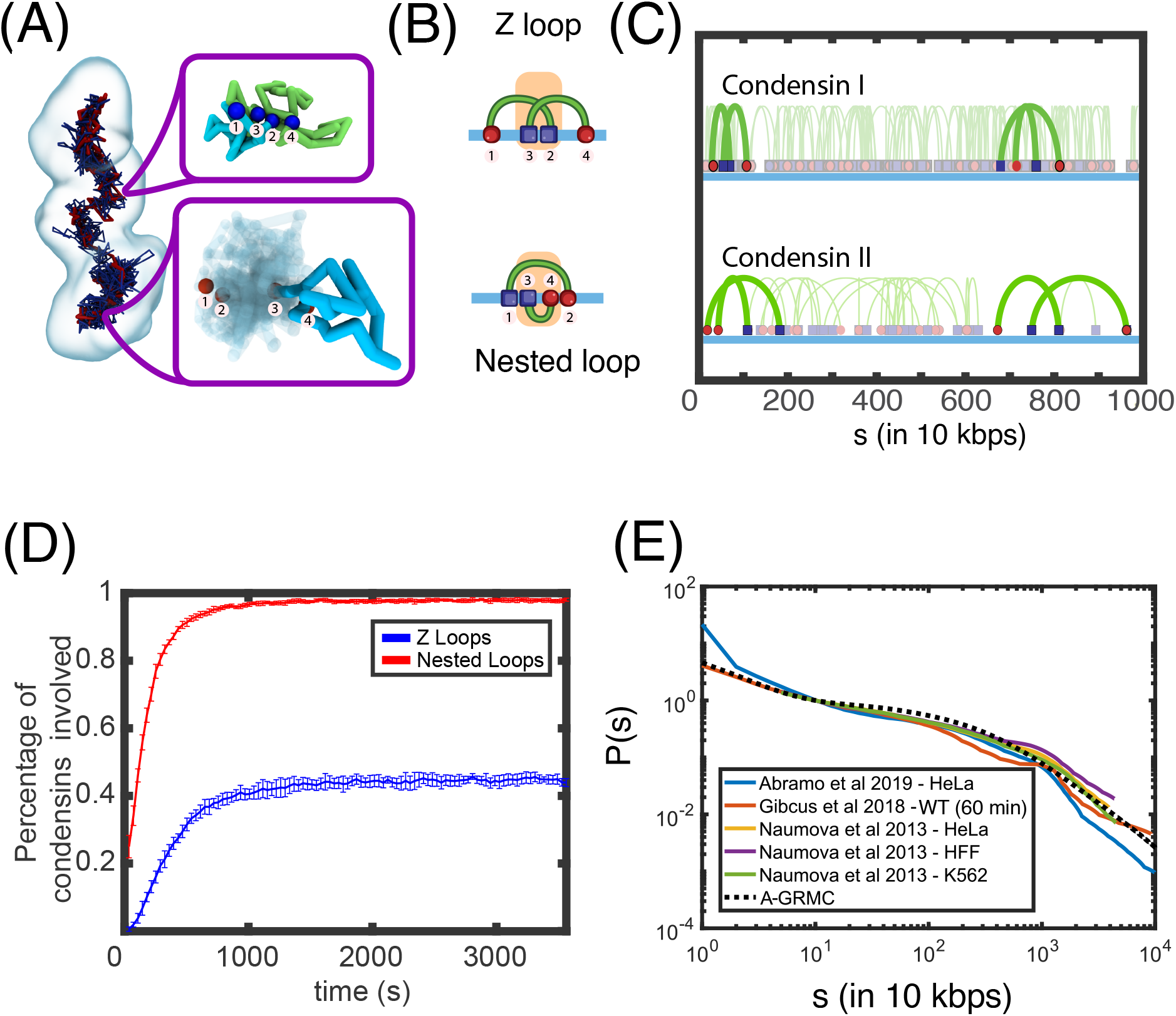
Loop generation mechanism. (A) A snapshot of a structure, generated in the A-GRMC simulations of the mitotic chromosomes, shows the backbone formed by the condensin molecules. Condensin I motors are shown in dark blue, whereas condensin II are in red. The magnified regions show an example of a Z-loop formed by two Condensin I molecules, and a nested loop formed by two Condensin II molecules respectively. For the Z-loop, the two loops are colored in cyan and green. For the nested loop the larger loop is shown in transparent colors, while the smaller loop is cyan. (B) Cartoon depicting two condensin molecules involved in a Z-loop and a nested loop. The overlapping segment of the chromatin between each loop is highlighted. (C) A snapshot of condensin I and II loops in 10 Mbps of DNA. Examples of both the Z-loops and the nested loops in either condensins are highlighted. (D) Fraction of condensin motors involved in the formation of a Z-loop or nested loop as a function of time. (E) *P*(*s*) in the metaphase chromosomes from different experiments and cell-types along with the predictions of the A-GRMC calculations (dashed lines). The A-GRMC calculations fall within the spread of the *P*(*s*) in the experiments.

### Symmetric loop extrusion (LE)

Ganji et al. 2018 have suggested that LE by a single yeast condensin motor is strictly asymmetric. In contrast, a more recent study (Kong et al. 2020) claims that LE by human condensin could be both symmetric and asymmetric. Both the mechanisms might be operative in *Xenopus* eggs (Golfier et al. 2020). It is possible that LE is mostly asymmetric during metaphase and symmetric during interphase (Banigan and Mirny 2020). In order to reconcile, at least partially, these seemingly opposing conclusions, we modified the LE scheme in order to investigate the consequences of symmetric LE on the *P*(*s*) as a function of *s*. As shown in Fig. 3(A), we exchanged the mobile and static sites randomly to allow the loop to form and grow on both sides. This is achieved by randomly switching the mobile and static sites with each other. If the probability of switching is equal, a loop could grow on both sides, thus creating symmetric extrusion. In Figs. 3 (B) and (C), we show two possible mechanisms for symmetric LE within the scrunching mechanism. Fig. 3 (B) illustrates that the loop itself is held between the two heads, while the hinge randomly binds to either side of the loop, resulting in symmetric LE. In Fig. 3 (C), the loop is held at one of the two heads, and the hinge binds and reels in DNA from the opposite side. In this scenario, condensin has to shuttle the loop back and forth between the two heads, thus creating an effective symmetric extrusion. Experiments could be designed to assess the validity of either of the mechanisms. We find that both fully symmetric extrusion and asymmetric extrusion result in the same *P*(*s*) versus *s* (Fig. 3(D)).

**Figure 3:**
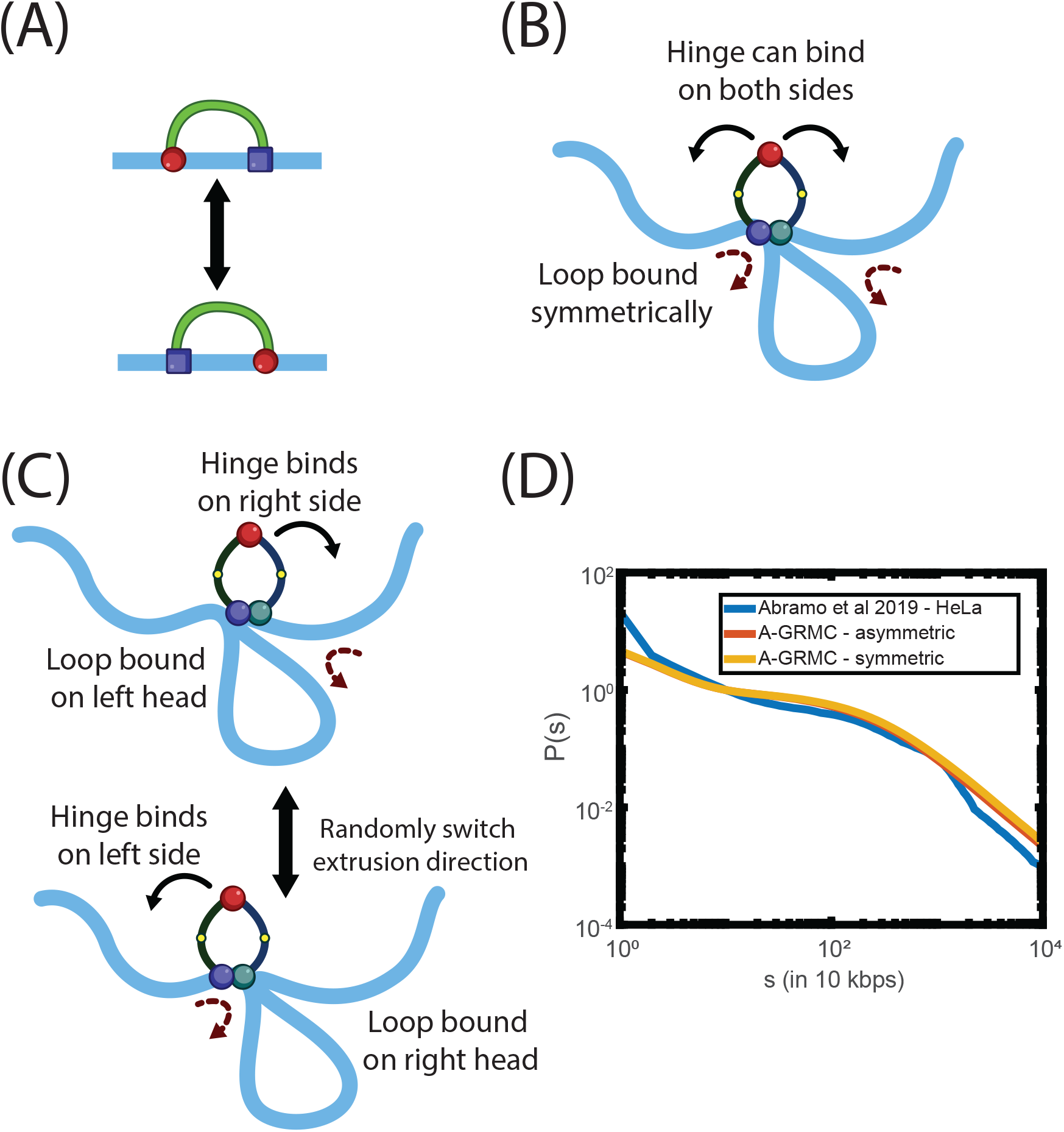
Symmetric versus asymmetric loop extrusion. (A) Symmetric extrusion is realized by exchanging the mobile and static sites (see also Fig. 1). Cartoon showing possible mechanisms of symmetric extrusion according to the scrunching mechanism. (B) The hinge can bind to both sides of the loop and the loop is held between the two heads. (C) The extrusion switches between two forms in the asymmetric extrusion. (D) Data for *P*(*s*) versus *s* for mitotic chromosomes for HeLa cells from experiments Abramo et al. 2019 compared with A-GRMC calculations using both symmetric and asymmetric extrusion. Clearly both the symmetric and asymmetric extrusion can reproduce the same *P*(*s*) curves.

### HeLa Cells

Interestingly, Fig. 3(D) shows that asymmetric extrusion coupled with the natural formation of *Z*-loops and nested loops also reproduces mitotic chromosome structures in HeLa cells (Abramo et al. 2019). A 100 Mbps of HeLa cell chromosome has ≈ 343 condensin II and ≈ 2, 553 condensin I molecules, which were calculated based on experimental measurements (see the Method section and the SI). Out of these, on an average ≈ 210 condensin II and ≈ 1205 condensin I molecules are bound to the chromosome. We found that, out of 1415 bound condensin molecules, ≈ 97% are involved in one or more nested loops, and ≈ 45% participate in one or more *Z*-loops. These results show that overlapping loops and *Z*-loops ensure that the loops generated by LE by multiple motors densely cover the entire chromosome without gaps.

### Time-dependent changes in the mean loop length

The dependence of the average loop size changes as a function of time. Fig. 4(A) shows that the loop sizes, averaged over 500 LE realizations, increase rapidly and saturate to a fixed value. The individual traces of the average loop sizes for five realizations reveal little change from one initial condition to another (Fig. 4(B)). The average size of the loops generated by condensin I (≈ 250 kbps) is about six times smaller (Fig. 4(A) than those generated by condensin II (≈ 1, 500 kbps). The long-range contacts established by condensin II motors likely stabilizes the helical scaffold where the multitude of short-range contacts, driven by condensin I, are responsible for overall compaction. Similarly, it takes longer (by about an order of magnitude) for condensin II to extrude a longer loop, with the LE velocity being the same.

**Figure 4:**
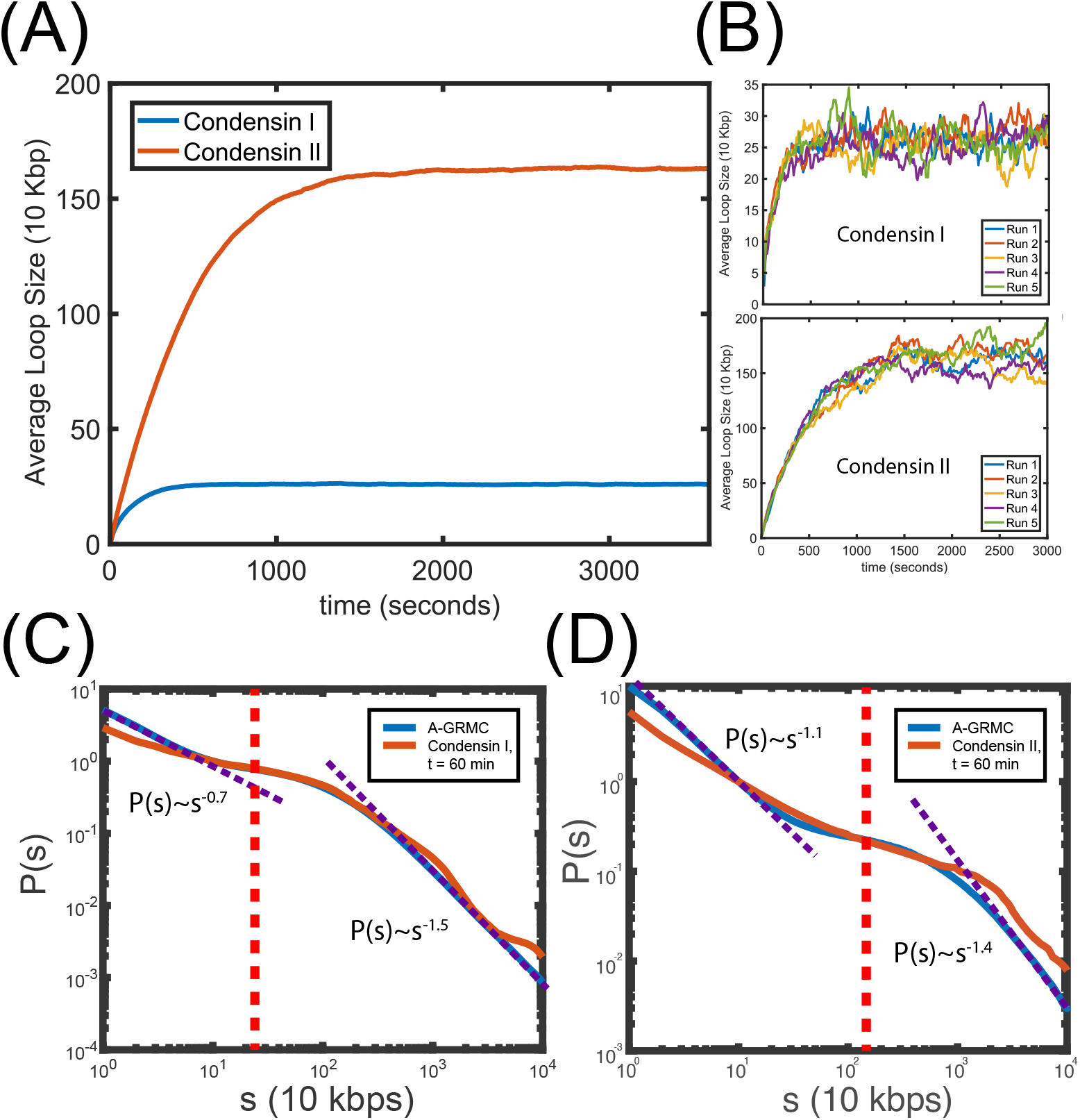
Time evolution of loop-sizes and *P*(*s*) curves for DT40 cells. (A) Average loop size as function of time for condensin I without condension II (blue) and condensin II without condensin I (red) from A-GRMC calculations. The mean loop sizes generated by condensin II are greater than the ones generated by condensin I even though the number of condensin I is larger than condensin II. (B) Evolution of average loop size in 5 independent simulations for condensin I and II. The figure shows that the mean loop size is reached rapidly within 1,000 seconds after the extrusion process starts. (C) Comparison of the dependence of *P*(*s*) on *s* predicted by A-GRMC calculations and experiments at *t* = 60 min upon depletion of condensin II. (D) *P*(*s*) versus *s* for condensin I depleted chromosomes at *t* = 60 min compared with the A-GRMC calculations. The red-dotted line indicates the average loop-length. Both (C) and (D) show a much sharper decrease in *P*(*s*) when *s* exceeds the average loop length. The small and larger exponent values are calculated between 1-90 Kbps and 10-100 Mbps, respectively.

Comparison of the calculated *P*(*s*) as a function of *s* using simulations and cells without condensin II at *t*=60 min (Fig. 4(C)) shows reasonable agreement. The decay of *P*(*s*) occurs in two steps. On the genomic length, *s* ≤ 1*Mbp*s, coinciding roughly with the size of the TADs, we find that *P*(*s*) ∼ *s*^*−*0.7^ whereas *P*(*s*) ∼ *s*^*−*3*/*2^ for *s >* 1Mbps. The values of the two exponents, calculated using the A-GRMC theory, are in reasonable agreement with experiments, especially considering that the errors in the measurements are not easy to estimate. Fits to the experimental *P*(*s*) versus *s* (Fig. 4(C)) yield *P*(*s*) ∼ *s*^*−*0.5^ (*s* ≤ 1*Mbp*s) and *P*(*s*) ∼ *s*^*−*1.9^ (*s >* 1Mbps). Although qualitatively similar, the agreement between the calculations and experiments for *P*(*s*) in condensin I depleted cells is less satisfactory for *s* ≤ 1*Mbp*s (Fig. 4(D)). The calculated exponent and that extracted from the Hi-C data are 1.1 and 0.7 respectively. On scales greater than *s >* 1Mbps the values of the exponent characterizing the decay of *P*(*s*) coincide.

### Effects of condensin I and condensin II depletion

Since both condensin I and II contribute to the mitotic chromosome formation, it is important to determine the differential contribution of the two motors in determining the dependence of *P*(*s*) on *s*. It is known that condensin I and condensin II bind to chromosomes independently (Hirota et al. 2004; Ono, Losada, et al. 2003), with the latter extruding larger loops due to the increased lifetime of the bound state (Fig. 4(A)). The distinct roles played by condensin I and condensin II were examined by selectively depleting the two motors(Gibcus et al. 2018). It was noticed that when both the motors are simultaneously present, there is a peak in the *P*(*s*) in the DT40 chromosomes from the DT40. The location of the peak moves to larger genomic distances (*s* values increase) accompanied by an increase in the magnitude of *P*(*s*), as the cell cycle proceeds towards mitosis. In order to determine the origin of the bump, the motors were selectively degraded, allowing them to infer *P*(*s*) as a function of *s* in the presence of only one of the motors. An important experimental finding is that the expected helical structure in mitotic chromosomes is predominantly driven by condensin II, with condensin I playing little or no role. In order to assess the accuracy of the A-GRMC theory, we calculated *P*(*s*) by selectively depleting the condensin motors.

The results for condension II depleted DT40 cells are shown in Fig. 5A, Fig. 5C, and Fig. 5E. Using *P*_*step*_ equal to unity produces reasonable agreement between the calculated and measured *P*(*s*) as a function of *s*. We optimized the step size to obtain the smallest error between simulations and experiment. The values of the optimized *P*_*step*_ do change with time (Fig. 5A), which implies that as the mean step size that the motors take vary as mitosis progresses. The optimal values of *P*_*step*_ as a function of time are displayed in Fig. 5C. Fig. 5(E) shows that, at the optimized *P*_*step*_, our theory quantitatively reproduces the measured (Gibcus et al. 2018) *P*(*s*) curves at all times. This suggests that the proposed mechanism for extruding loops by multiple condensin I motors has some validity. Based on the results in Fig. 5 we speculate that the variation in *P*_step_ implies that the mechanical tension along the chromosomes changes as cell progresses to mitosis.

**Figure 5:**
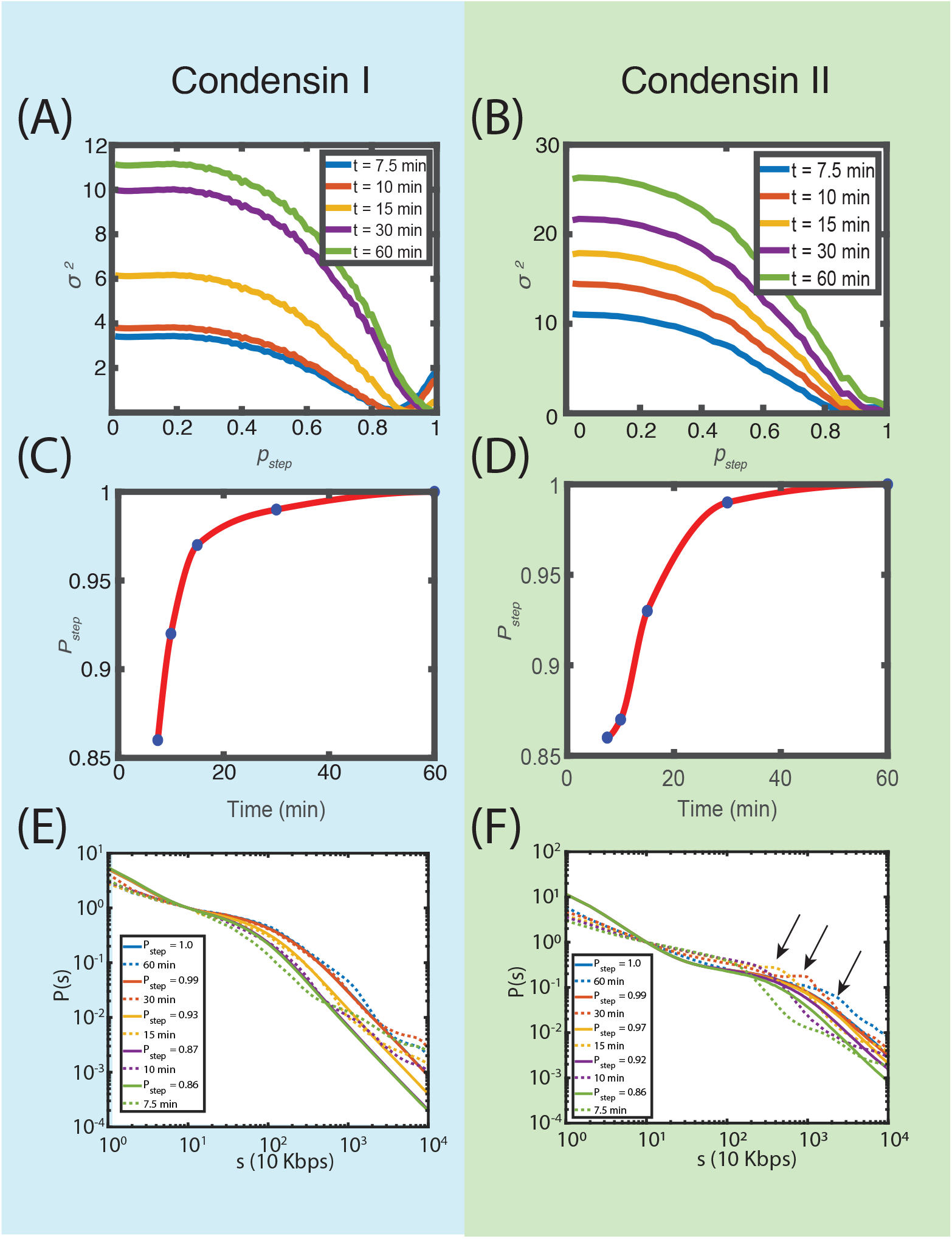
Effect of step size on *P*(*s*) for DT40 cells. Panels A, C, and E are for condensin II depleted cells ans B, D, and F are the results for condensin I depleted cells. (A) Mean-squared error (*σ*^2^, defined in Eq. (15) in the SI) between *P*(*s*) calculated using the A-GRMC and Hi-C at different times for condensin I. (B) Same as (A) except these are for condensin I depleted cells. (C) Optimal values of *P*_*step*_ corresponding to the *P*_*step*_ that produces the minimum in *σ*^2^ in (A). The lines are from theory and the dashed curves are experimental data. (D) Same as (C) except the results for condensin I depleted cells. (E) Comparison of *P*(*s*) versus *s* at different times between the theoretical predictions and experiment for condensin II depleted cells. The optimal values of *P*_*step*_ are given. (F) Same as (E) except the results are for condensin I depleted cells. The arrows indicate the bumps in the *P*(*s*).

The results for condensin I depleted cells are shown in Fig. 5B, Fig. 5D, and Fig. 5F. The results for *P*(*s*) as a function of *s* calculated at the optimum *P*_*step*_ values (Fig. 5D; the errors between the calculated and experimental *P*(*s*) are in Fig. 5B). Although the agreement between theory and experiments is reasonable (Fig. 5F), the simulations do not capture the bump in the *P*(*s*) when only condensin II is present. The bump in *P*(*s*) is an indication of the underlying helical structure of mitotic chromosome (Gibcus et al. 2018). In order to obtain helical structures, the model must contain anisotropic interactions involving the chromosomes and the motors. The absence of such interactions in the A-GRMC, which is likely generated by condensin II by an unknown mechanism, explains the lack of the pronounced bump in *P*(*s*) when condensin I is depleted.

### Impact of the motor residence time, the pausing time, and the density of motors on *P*(*s*)

We explored the effects of changing the parameters in the A-GRMC on *P*(*s*) as a function of *s*. The results are shown in Fig. S4 in the SI.

#### Life time of bound condensin motors

Fig. S4(A) in the SI shows that increasing the residence time of condensin motors leads to additional long-range contacts, which is reflected in a plateau-like feature in *P*(*s*) for *s >* 10^5^ bps. As shown in Eq. 1 in the SI, the unbinding probability of condensin molecules, *p*_off_, is inversely proportional to the residence time. Thus, increasing the residence time decreases *p*_off_, which in turn increases the lifetime of condensins that are bound to the DNA. As a consequence, longer loops form, which enhance the *P*(*s*) value at large *s*. A corollary of this finding is that the longer loops driven by condensin II is due to the enhanced residence time of condensin II (≈ 6 min) on the chromosome compared to condensin I (≈ 2 min).

#### Pausing time

The effect of changing the pausing time, *τ*_*p*_, has the opposite effect compared to increasing the residence time. Fig. S4(B) in the SI shows that upon increasing *τ*_*p*_ the probability of contact formation for *s >* 10^5^ bps sharply decreases. Comparison of the predicted and measured *P*(*s*) shows that optimal value of *τ*_*p*_ is about 4 ∼ 8 seconds. Our theory, therefore, could be used to extract the residence and the pausing times of condensin motors by fitting the predictions to the experimental *P*(*s*) versus *s*.

### Number of bound motors

The number of bound condensins (*n*_*b*_) also influences the shape of the *P*(*s*) curves, as revealed in Fig. S4(C) the SI. Because *p*_on_ ∝ 1*/*(*n*_*t*_*/n*_*b*_ − 1) (see Eq. 2 in the SI), it follows that increasing *n*_*b*_ while keeping *n*_*t*_ constant leads to larger binding probability *p*_on_. The increase in the density of bound condensins to DNA further results in more loops being extruded. In addition, the values of *P*(*s*) should be larger at large *s* values.

However, once all the available condensins are bound, the *P*(*s*) would not change. The best fit to experiments shows that on an average the number of bound condensin I is *N*_*b*_ = 255 (Fig. S4(C) in the SI). We also calculated *P*(*s*) by varying the *p*_on_*/p*_off_, Fig. S4(D), which is similar to changing *n*_*b*_*/*(*n*_*t*_ − *n*_*b*_). Higher values of *p*_on_*/p*_off_ results in more condensins being bound to the DNA, resulting in the extrusion of longer loops.

Taken together, these results suggest that the biologically relevant parameters *τ*_*p*_, *τ*_*B*_ and *n*_*b*_ are optimized to extrude loops of appropriate sizes. It is likely that these parameters depend on the chromosomes and the cell type as well.

### Helical structures of mitotic chromosomes are predicted using a data-driven method

To assess if the bump in *P*(*s*) could be reproduced, and explained in terms of helical (or related) conformations, we used the recently developed HIPPS method (Shi and Thirumalai 2021). As a complement to the A-GRMC simulations and to compare the extent of helix formation in DT40 and HeLa cells, we calculated the *P*(*s*) at different time points as the cell transitions from the interphase to the mitotic phase using the HIPPS method, which predicts an ensemble of 3D chromosome structures with the measured Hi-C maps as the only input (see Methods).

The *P*(*s*) as a function of *s*, calculated using the 3D structures (Fig. 6) shows that the calculated contact probability for condensin I depleted chromosomes in DT40 cells (Fig. 6A) and experiments are in *quantitative* agreement at all times. In particular, the HIPPS method accounts for the pronounced peak (Fig. 6B), which we show below is important in producing structures with random helix perversion (RHP). Naturally, the predictions are also quantitative for DT40 chromosomes when both the motors are present (Fig. 6C). It is also gratifying that the calculated and measured *P*(*s*) for HeLa cells is also accurate (Fig. 6D).

**Figure 6:**
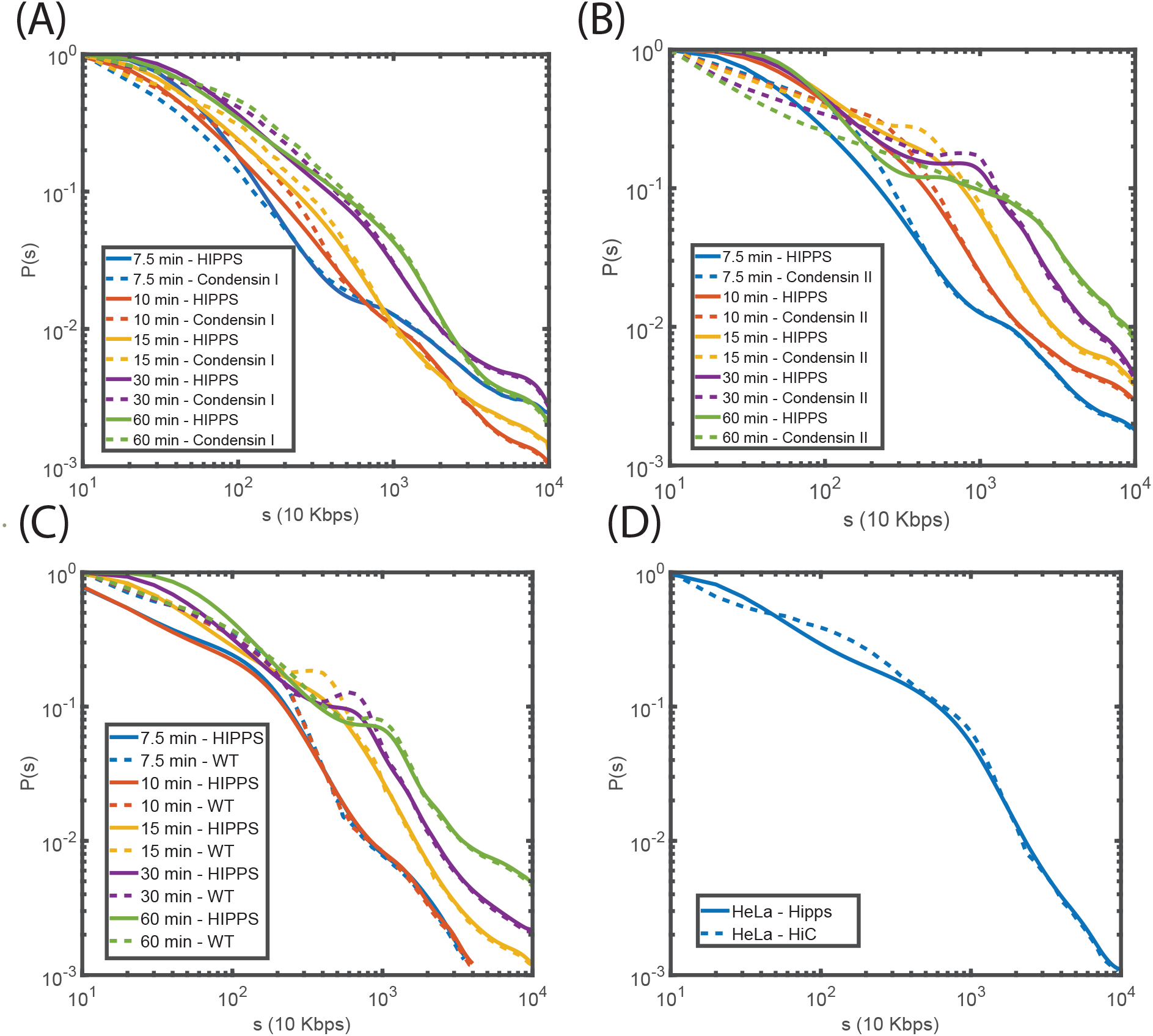
Comparison of calculated *P*(*s*) using the HIPPS method (solid lines) with experiments (dashed lines). (A) *P*(*s*) from experiments for DT40 cells with only condensin I (condensin II depleted) at various times compared with the HIPPS calculations. Same as (A) except the comparison between the HIPPS predictions and experiments correspond to condensin I depleted cells. (C) [*P*(*s*), *s*] from experiments for cells with both condensins (wild type) at various times compared with the corresponding HIPPS calculations. The results in (A), (B), and (C) are for DT40 cells. (D) Comparison between HIPPS predictions and experiments for Hela cells Abramo et al. 2019.

### Periodicity in angular correlation functions

With the 3D coordinates of the loci at hand, we quantified the difference in the extent of helix formation at different times in the DT40 cell, as it goes through the phases in the cell cycle, using the angular correlation function,

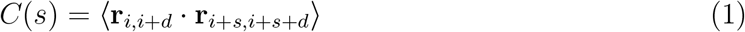

where **r**_*i,j*_ is the vector between the *i*^*th*^ and *j*^*th*^ loci and *d* = 32 (Shi and Thirumalai 2021). It can be shown that for a perfect helix, *C*(*s*) should be periodic with a period that is proportional to the helical pitch (Fig. S5 in SI). From Fig. 7(A), which shows *C*(*s*) as a function of *s* upon condensin II depletion at different times, we notice that there are small but discernible peaks at *t* = 15, 30 and 60 minutes. The locations of the peaks change from *s* ≈ 2.5 Mbps (*t* = 15 mins) to *s* ≈ 7.5 Mbps (*t* = 60 mins). Thus, even in the absence of condensin II, we predict that there is a tendency towards helix formation, which is likely masked in the isotropic ensemble-averaged *P*(*s*), especially if the bump in *P*(*s*) is not prominent. It is worth noting that both experiments and our calculations show that there is a minor shoulder in the *P*(*s*) plots even in the absence of condensin II (Fig. 6A).

**Figure 7:**
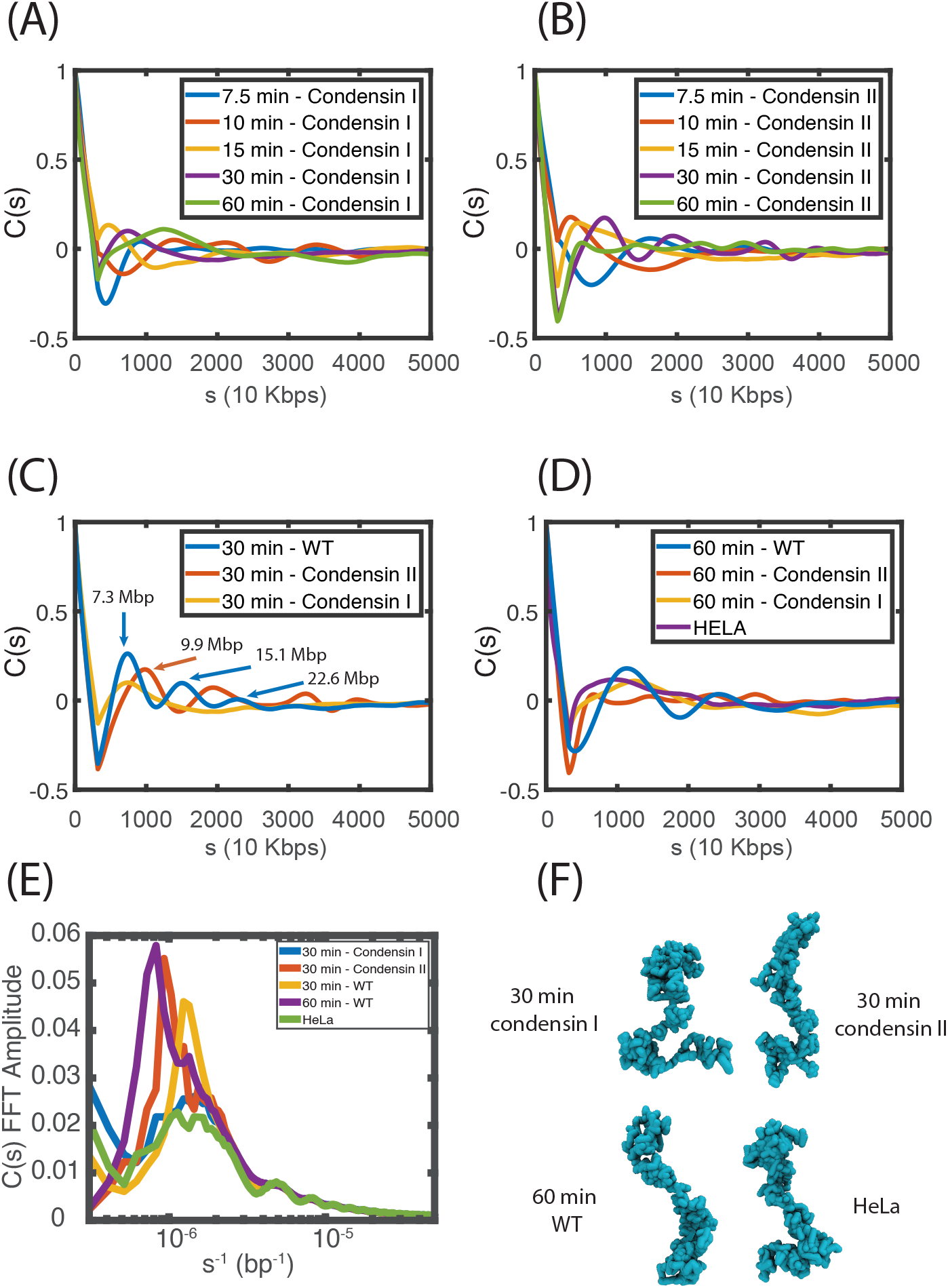
Angular correlation (*C*(*s*)) as a function of genomic distance at different times. (A) *C*(*s*) as a function of *s* at various times for DT40 cells without condensin II. A peak at *s* ≈ 5*Mb* with a small amplitude that is most pronounced at *t* = 15mins (orange line) suggests that condensin I alone induces helix formation but not as much as condensin II. *C*(*s*) as a function of time for condensin II chromosomes (condensin I depleted) DT40 cells. The periodicity (roughly 10 Mbs), especially at *t* = 30 min, is a signature of helix formation. (C) Comparison of *C*(*s*) versus *s* for WT(blue) condensin I depleted cells (orange), and condensin II depleted (red) DT40 cells at *t* = 30 min. (D) Same as (C) except these correspond to *t* = 60 min. Also shown is *P*(*s*) versus *s* for HeLa cells (violet), which illustrates less prominent helicity. (E) Fourier transform of (*C*(*s*)) as function of *s*^*−*1^ shows that in the amplitude spectrum condensin II chromosomes demonstrate stronger peaks compared to condensin I chromosomes. The peaks HeLa cells are weaker peaks compared to DT40 cells. (F) Example structures constructed from HIPPS for *t* = 30 mins condensin I/II along with WT DT40 cells at *t* = 60 mins and HeLa cells. The structures show that the HIPPS method accurately predicts the expected conformal features in the mitotic chromosome.

When only condensin II is present (CAP-H-mAID in the notation used in experiments), both the amplitudes and the number of peaks increase (Fig. 7(B)), as time progresses. Just as in experiments (see the right panel in Fig. 5A in the experimental study (Gibcus et al. 2018)), Fig. 7(B) shows that the peak is most prominent at *t* = 10 min (red), *t* = 15 min (orange) and *t* = 30 min (purple). The peak positions shift to larger values of *s* as the cell cycle progress towards mitosis. Remarkably, these features are in *quantitative* agreement with experiments, without having to adjust any parameter to fit the experimental data. We compare the *C*(*s*) plots at *t* = 30 mins (Fig. 7(C)) and *t* = 60 mins (Fig. 7(D)) in the WT, condensin I and condensin II depleted structures. At both the time points, the periodicity is more prominent for condensin II depleted chromosomes than it is when condensin I is depleted. In accord with experiments, the extent of helix formation appears to be less prominent at *t* = 60 mins, compared to *t* = 30 mins.

### Helix formation is weaker in HeLa cell chromosomes

It is instructive to compare the *s*-dependent *C*(*s*) for HeLa and DT40 cells (Fig. 7(D)). There are two significant differences: (i) The peak position in *P*(*s*) for the chromosomes in the HeLa cell (purple line in Fig. 7(D)) is at a smaller *s* value compared to the DT40 cells at *t* = 60 mins (blue in Fig. 7(D)) (ii) More importantly, the number of peaks, indicating the extent of helix propagation along the genomic distance, is less in the HeLa cell relative to the DT40 cell. This shows that the helix formation depends on the species and the cell type. Examples of the structural ensemble are given in Fig. 7F.

The amplitude of the Fourier transform of *C*(*s*) (Fig. 7E) for condesin I/II chromosomes at *t* = 30 and *t* = 60 minutes, for the wild-type DT40 and mitotic HeLa cells show prominent peaks for condesin II and DT40 WT cells, containing both the motors. In contrast, the amplitudes are smaller for HeLa cells and condensin II depleted DT40 cells, implying that the degree of helix structure is less. Examples from the structural ensemble are shown in Fig.7(F).

### Random helix perversion (RHP) in mitotic chromosomes

The precise structural features of mitotic chromosomes have not been firmly established even though several studies (Ohnuki 1965; Marsden and U. Laemmli 1979) have suggested that they are helical. That there is an underlying periodicity in the structures is clear from the *C*(*s*) as a function of *s* (see Fig.7 and Fig. S5) is another indication of helical conformations. However, from these results we cannot determine whether the helix is left or right handed because *C*(*s*) = *C*(−*s*)). For instance, a perfect helix *C*(*s*) exhibits identical behavior regardless of the handedness (See Fig. S5 in the SI). Similarly, simulations that impose helical conformations and adjust the parameters to fit the *P*(*s*) data also cannot determine the handedness unambiguously because *P*(*s*) does not contain the information required to make this distinction.

We used the 3D structures to characterize the nature of the mitotic structures. To get insights into plausible arrangements, we first created synthetic structures that are perfectly helical (left or right handed). Not unexpectedly, *C*(*s*) for these structures are perfectly periodic with peaks (and valleys) occurring at *s* values that are integer multiples of the helix pitch, *P*(measured in Mbps units) (see Fig. S5 in the SI). However, they do not match the calculated *C*(*s*) from the 3D structures determined using HIPPS. We next generated synthetic structures with RHP by randomly altering the handedness across the entire mitotic chromosomes (see the SI for details). The random feature is controlled by the parameter *r*, which is zero for a perfectly periodic helix. For *r* = 0.01, the *C*(*s*) for the synthetic structures and the results in Fig.7B are similar.

The results in Fig. S5 suggests that in all likelihood mitotic structures have random helix perversion (a word coined by Maxwell as reported elsewhere (Goriely and Tabor 1998; McMillen, Goriely, et al. 2002), with alterations in the handedness occurring randomly. To affirm this possibility, we calculated the helix order parameter (Moradi et al. 2009) expressed in terms of an angle *χ*, which could be used to extract the local handedness (see SI for definition of *χ* and the implementation in the context of mitotic chromosomes). Using the 3D coordinates of 1,000 mitotic structures at *t*=30 minutes, we calculated distributions of *χ* for the ensemble of structures and a single structure. The *χ* distribution, averaged over the 1,000 conformations is broad, with a mean *χ* ≈ 0. Although for a single conformation, the distribution of *χ* is peaked at *χ* /= 0, it is not far from zero. Indeed, the *χ* distribution in Fig. S6(F) shows that both left and right handed chiral order, in almost equal number, coexist in a single conformation. We conclude that each conformation, with signature of RHP globally, may only have modest propensity to be either left or right handed.

## Discussion

We developed a theory based on loop extrusion driven by multiple condensin I and condensin II motors to determine the structural changes that occur in chromosomes as the cell cycles from the G2 phase to metaphase. The resulting A-GRMC has two components: (a) a kinetic scheme to determine the dynamics of loop extrusions mediated by multiple condensin I/II molecules, (b) a polymer model which takes the location of loop anchors generated by the condensin motors as input to generate the three-dimensional structures of chromosomes. The kinetic scheme is rooted in the scrunching mechanism, discovered first in the context of DNA transcription bubble formation in bacteria (Kapanidis et al. 2006), which has recently been invoked to explain the LE mechanism of a single condensin motor (Ryu et al. 2020; Takaki et al. 2021). Using the A-GRMC together with the results from HIPPS method, we have quantitatively accounted for all the experimental data. (1) The A-GRMC simulations accurately predict the behaviour of *P*(*s*) for mitotic chromosomes in HeLa cells and DT40 chicken cells when condensin II is depleted. (ii) The predictions using the A-GRMC, are less accurate when only condensin II is present. In particular, the bump in the *P*(*s*), whose origin is linked to plausible helix formation in mitotic chromosomes, is not predicted by the A-GRMC theory. (3) Remarkably, the calculations using the HIPPS method predict, with high accuracy, the behavior of *P*(*s*) for DT40 cells in the presence of both the motors, and upon depletion of the two motors, one at a time. In a sense, the HIPPS method solves the problem of predicting helical or structures with periodicity as the cell cycle progresses from the interphase to mitosis. (4) We found that the extent of helix formation in mitotic chromosomes depends on the cell type. For the HeLa cells, the peak in *P*(*s*) is less pronounced compared to DT40, which is also reflected in the angular correlation function (*C*(*s*)). (5) We find that even with only condensin I present, there are signatures of weak helix formation in *C*(*s*) calculated from the 3D structures. Interestingly, the structures of the mitotic chromosome have the characteristics of random helix perversion in which the helix handedness changes randomly throughout the chromosomes.

### Dynamical nature of the chromosomal axis

The cylinder shaped mitotic chromosome structure may be thought of having a scaffold axis, which traces the positions of both condensin I and I. The extruded loops, with variable length, are stapled to the scaffold. Because the mitotic chromosome axis is pictured to be helical, the loops that emanate from the motor proteins trace a helical path. The path traced by the motors are not static because their formation is controlled minimally by three time scales that determine the dynamical nature of the helical axis. (i) One is the average LE time. Using the mean loop lengths for condensin I (≈ 25 kbps) and 150 kbps for condensin II (Fig. 4A) and the LE velocity (≈ 1,500bps/s) for extrusion by a single motor, the extrusion time is 200 s for condensin I and 1,000 s for condensin II. When multiple motors are involved, the time increases by an order of magnitude (Fig. 4(A)). (2) The life times for bound condensins are estimated to be 120 s and 360 s for condensin I and condensin II, respectively. (iii) The overall transition time to go from the G2 phase to prometaphase is about 30 minutes in DT40 cells. The first two time scales are less than the cell cycle time, which implies that the motors attach and detach multiple times as the cells transition from the G2 to prometaphase. This implies that the loops must be dynamic, and more importantly, the scaffold is unlikely to be static (Gibcus et al. 2018). Only by averaging over many realizations does an ensemble-averaged backbone structure, with possibly complicated helical arrangement (see below), emerges.

### Arrangement of loops

Microscopy experiments (K. Samejima et al. 2012; Adolph et al. 1977) have suggested that the shape of the chromosome is cylindrical with the condensin and topoisomerase molecules forming the central axis. It is thought that the loops are roughly perpendicular to the putative central axis, which is dynamic as evidenced by the observed heterogeneity of the structures. Our calculations show that the loops are not arranged in a strictly consecutive non-overlapping fashion because the orientations and the binding sites of the condensins are random, and vary from one realization to another. The combination of nested and Z-loops creates structures that bear some resemblance to polymer bottle brushes, except that the central axis dynamically changes because the motors bind and unbind several times during the cell cycle.

### Roles of condensin I and condensin II in shaping the mitotic chromosomes

Since the pioneering discovery of condensin II (Ono, Losada, et al. 2003), there have been several studies (Hirota et al. 2004; Ono, Fang, et al. 2004a; Hirano 2016) that have probed the differing roles these motors play in the formation of mitotic chromosomes. Although condensin I and II load onto chromosomes independently, with the former gaining access only after breakdown of the nuclear envelope, both are required for mitotic chromosome assembly. It has been suggested that (Green et al. 2012) condensin II imparts rigidity to the chromosomal axis whereas condensin I enables lateral compaction.

The A-GRMC simulations show that condensin II extrudes longer (by a factor of six) compared to condensin I. This implies that the establishment of longer-range contacts between distant loci requires condensin II whereas condensin I assists in the generation of shorter ranged contacts. Coincidentally, the short-range contacts occur on length scales that are characteristic of TADs, whereas the genomic size of long-range contacts is associated with compartment sizes. The 3D structures generated by the HIPPS method shows (Fig.7B) an emergent length scale at *s* ≈ 10 Mbps in *C*(*s*), which coincides with the peak in *P*(*s*) at *t* = 30mins (see Fig. 6B). To the extent rigidity arises from the periodic structure, we could surmise that condensin II might determine the backbone stiffness (Hirota et al. 2004).

### Random helix perversion

Although the radial loop model, extracted from micrographs (Marsden and U. Laemmli 1979), seem to suggest that mitotic chromosomes are left handed, this issue has not been settled, and has come to focus in light of a recent imaging study of mitotic chromosomes (Chu et al. 2020). They propose that the helix is best described as having, PHP, periodic helix (or tendril) perversion, a structure that was first noted sometime ago in creeping plants (Darwin 1875). The PHP is a perfect repeat of monomer units. Each unit consists of one left and one right handed helix. For structures with PHP, *C*(*s*), is shown in Fig. S5 for synthetically generated structures. Comparison with the *C*(*s*) predicted by the HIPPS method and *C*(*s*) for structures with PHP shows qualitative differences. Although we have established that the HIPPS method that uses Hi-C data as input, quantitatively reproduces *P*(*s*) at all time points, the consequences of ensemble averaging and lack of error estimates in the Hi-C experiments make it hard to assess the reliability of drawing firm conclusions about the details of mitotic structures. Therefore, additional super resolution imaging studies, which can determine structures at the single chromosome level, will be required to definitely address the nature of helix perversion in mitotic chromosomes.

The conclusions reached here concerning RHP are most consistent with a recent study (Zhang and Wolynes 2016), which used an entirely different approach. In that study, the Hi-C data for mitotic chromosomes was used as input to determine the parameters of an assumed energy function using the maximum entropy method. Most interestingly, they found that the structures, obtained by simulations of the energy functions with learned parameters obtained by fitting to the Hi-C data, showed that the overall shape of the mitotic chromosomes is cylindrical, consisting of a mixture of left and right handed helices implying that chiral symmetry is spontaneously broken. The resulting structures on an average are likely to be similar to our findings.

In summary, we have introduced a model, which could be a reasonable starting point for elucidating the mechanism for predicting multiple condensin-driven structural changes that occur during the mitotic cycle. In combination with the HIPPS method, we have derived plausible structures of mitotic chromosomes. Despite some success, the difficult problem of cooperation between hundreds of condensin (and other) motors is a daunting challenge.

### Limitations of the study

The most glaring weakness of our study is that the calculated *P*(*s*) using the A-GRMC model in the presence of only condension II (condension I is depleted) is not in near quantitative agreement with experimental measurements. The chromosome structures, upon condensin I depletion, does have a cylindrical shape but the bump in *P*(*s*) that moves to larger values of *s*, as the cell cycle proceeds towards mitosis, is not captured. Although we quantitatively reproduced the experimental *P*(*s*) using the HIPPS method, it does not reveal the mechanism by which multiple condensin II motors extrude long loops to generate the RHP structures. The differences in the *P*(*s*) obtained upon depletion of condensin I (condensin II) suggests that there could be subtle but important difference in the loop extrusion mechanism employed by the two motors even though their structures, except for the non-SMC parts, are similar.

## Methods

We developed the Active GRMC (A-GRMC) model, which incorporates the collective *active* dynamics of multiple condensin motors in concert with the GRMC model (Bryngelson and Thirumalai 1996; Shi and Thirumalai 2019).The calculations using the A-GRMC are done in two steps. First, we perform kinetic simulations of loop extrusion (LE) driven by multiple condensin motors to determine the genomic location of loop anchors as a function of time. The motors are the anchor points to which the loops are stapled. Second, the dynamics of LE is used in conjunction with the GRMC to generate chromosome structures, which allowed us to calculate *P*(*s*) that can be compared with experimental data. The technical details are given below and further elaborated in the Supplementary Information (SI).

### Loop extrusions (LE) by multiple condensins

The model for multi-condensin loop extrusion is shown schematically in Fig. 1. Each condensin complex, belonging to the Structural Maintenance Complex, is modeled as a pair of capture points on the DNA. In accord with the scrunching mechanism (Takaki et al. 2021; Ryu et al. 2020), we assume that only one of the capture points (depicted as a circle in Fig. 1) moves along the DNA while the other point (depicted as a square) is fixed. In other words, each motor performs one-sided loop extrusion. Condensins disengage from the DNA track stochastically with probability, *p*_off_, or load onto DNA with probability *p*_on_. The direction of loop extrusion along the DNA is random and the mobile point moves processively (takes many steps before disengaging) along the direction of initial orientation. The system is initially (*t* = 0 min) in the prometaphase. At later times, the movement of the mobile-point is controlled by the value of the stepping probability, *P*_step_ (see SI for details). The values of *p*_on_ and *p*_off_ are calculated from number of bound condensins (*n*_*b*_) and total number of condensins (*n*_*t*_). These quantities have been measured in experiments (see Supplementary Information (SI) for details).

### Kinetic simulations for multi-condensin LE

The kinetic simulations for multi-condensin driven LE is performed in four steps (Fig.1): (1) At *t* = 0 min, the condensin motors are assumed to be in the unbound state. At each time step, individual motor (condensin I or II) binds to the genome with probability *p*_on_. A bound condensin can also disassociate with probability *p*_off_. (2) Once the binding and unbinding events are determined, each newly bound condensin is assigned a random orientation for LE, which is unchanged until the condensin disengages from the genome. (3) A bound condensin takes a single LE step with probability, *P*_step_, which is set to unity in all the metaphase chromosome calculations, unless specified otherwise. The location of the mobile point after a single LE step is determined by the step size of condensin, which is sampled from the distribution given in the SI (Eq. S3). If the motor encounters another condensin, it pauses (does not take any steps) for a time duration, *τ*_*p*_. (4) After executing these events, the clock is advanced by one time-step, and the first three steps are repeated. The parameters in the kinetic simulations, listed in Table 1, are determined using the experimental data. The details are in the SI.

The locations of the condensin heads constitute the scaffold from which loops of various sizes emanate. At each time step, the genome locations of the condensin heads are determined and used in conjunction with GRMC to obtain the chromatin structures.

### A-GRMC

The energy function for A-GRMC (Eq. S5 in the SI) has two terms: 1) the chain connectivity, modeled using harmonic bonds, with a spring constant, *k* = 3*k*_*B*_*T/b*^2^, where b is the locus size, 2) the looping interaction, which is also modeled using harmonic potentials between the two heads of condensin with the same spring constant, as given above. The dynamics of the loops are determined by the kinetic simulation described above and illustrated in Fig. 1. The loop anchors that are pinned by the motors serve as the backbone scaffold for the chromosome. At each time-step in the kinetic simulation, the loop anchors are fixed and their positions are known. Thus, the energy function (Eq. S5) is fully determined. Given the energy function, the chromosome structures can be generated (see details in the SI) and quantities of interest, such as *P*(*s*) as a function of *s*, can then be readily calculated (see details in the SI). Without loss of generality, we use reduced units by setting *k*_*B*_*T* = 1, and the locus size *b* = 1, which is on the order of a hundred microns in real units Shi and Thirumalai 2021.

Each coarse-grained monomer in the A-GRMC represents 10 kbps, which is longer than the persistence length of DNA and likely also the chromatin fiber (Sanborn et al. 2015), justifying the use of the flexible polymer model without any angle potential (Eq. S5). Although the fundamental length scale of the loop-extrusion step is comparable to the persistence length of DNA, the total chromosome length is orders of magnitude larger. Therefore, even though the step-size distribution is calculated using the Eq. S3 in the SI, which is derived for semi-flexible polymers, we model the chromosomes as a generalized Rouse polymer.

### Hi-C-polymer-physics-structures (HIPPS)

To complement the A-GRMC model for predicting the *P*(*s*) as a function of *s* throughout mitosis of in the DT40 and HeLa cells, we used the HIPPS method (Shi and Thirumalai 2021), whose sole input is the measured comtact maps from Hi-C experiments. Because the details of the HIPPS method, including the theoretical basis, and applications are given elsewhere (Shi and Thirumalai 2021; Shi and Thirumalai 2022), here we only provide a brief description. The algorithm implementing HIPPS has three steps: (1) First, the mean contact probability between two loci *i* and *j* is used to calculate the average spatial distance between them using polymer physics concepts (Shi and Thirumalai 2021). In practice, we produced smooth contact maps, *C*_*i,j*_ from *P*(*s*) inferred from the Hi-C experiments. The matrix *C*_*i,j*_ is then transformed to a distance map. (2) The mean distance matrix between all the loci is used as a constraint to construct the probability distribution of the loci coordinates using the maximum entropy principle. The HIPPS method generates an ensemble of 3D coordinates for the loci using experimental Hi-C map *without* constructing any polymer model. The *P*(*s*) versus *s* plots can be readily calculated from the 3D coordinates. Shi and Thirumalai 2021 showed that the HIPPS method is remarkably successful in predicting the 3D structures of all the 23 interphase chromosomes from GM12878 cells. The code to implement the HIPPS method can be accessed at GitHub.

## Acknowledgements

We thank Kiran Kumari, Xin Li, and Sucheol Shin for useful discussions. This work was supported by a grant from the National Science Foundation (CHE19-00033) and the Welch Foundation through the Collie-Welch Chair (F-0019).

