## Supplementary Information for "Structural changes in chromosomes driven by multiple condensin motors during mitosis"

*Department of Chemistry,*

### 1 Estimates of $p_{on}$ and $p_{off}$

The theory for generating the shape of the mitotic chromosomes is based on the model described in Fig. 1 in the main text. Two key parameters needed for the simulation of the Active-Generalized Rouse Model for Chromosomes (A-GRMC) are, $p_{on}$ and $p_{off}$, which are the probabilities that condensin motors bind and unbind from DNA, respectively. We calculated $p_{on}$ and $p_{off}$ using data from Fluorescence Recovery After Photo (FRAP) experiments (Walther et al. 2018) for the residence times of condensin I and II bound to chromosomes. Since the mechanisms of the binding-unbinding processes are unknown, we assume that they obey first order kinetics. We assumed that at each step in the simulations condensin binds or unbinds with probability $p_{on}$ and $p_{off}$, respectively. The average time (in units of the a time step = 1 sec - see the next section for an explanation) a single condensin stays bound to a chromosome is,

$$\langle \tau_B \rangle = \sum_{t=1}^{\infty} t \cdot p_{off} \cdot (1 - p_{off})^{(t-1)} = \frac{1}{p_{off}}. \tag{1}$$

Since the number of bound ($n_b$) and unbound ($n_u$) condensin molecules is relatively constant during mitosis (Walther et al. 2018), we assume that they are in equilibrium with each other. Given that the total number of condensins $n_t = n_b + n_u$, it follows that at equilibirum,

$$n_u \cdot p_{on} = n_b \cdot p_{off} \implies p_{on} = \frac{n_b \cdot p_{off}}{n_u}. \tag{2}$$

Using the data in Tables S1 and S2, we calculated $n_t$ for both the condensin motors. Walther et al. 2018 used diploid HeLa cells whose genome size is $N = 2$ x 7.9 Gbp. Assuming uniform coverage of condensins in 100 Mbps of DNA, $n_t = 403,395 \cdot 100 \cdot 10^6/(2 \cdot 7.9 \cdot 10^9) \approx 2553$ for condensin I and for 100 Mbp of DNA $n_t = 54,219 \cdot 100 \cdot 10^6/(2 \cdot 7.9 \cdot 10^9) \approx 343$ Condensin II. Because $n_b$ for condensin I varies during mitosis, we used the highest value of $n_b$ which is formed during prometaphase (shown in Table S3). Thus, for a 100 Mbps DNA $n_b = 190433 \cdot 100 \cdot 10^6/(2 \cdot 7.9 \cdot 10^9) \approx 1205$ for condensin I. The total number of condensin II molecules remain relatively constant, so we used the average number from Table S4, which leads to $n_b = 33180 \cdot 100 \cdot 10^6/(2 \cdot 7.9 \cdot 10^9) \approx 210$.

We used the average value of bound time $\langle \tau_B \rangle = 120$ s, for condensin I and $\langle \tau_B \rangle = 360$ s, for condensin II. The use of average values did not significantly change any of the results. Using these values in Eqs. 1 and 2, we calculated $p_{on}$ and $p_{off}$ for condensin I and II.

The parameters, listed in the Table 1 in the main text, show that $p_{on}$ and $p_{off}$ are different for condensin I and II. All the six parameters needed in the A-GRMC simulations are obtained from experiments. It is in this sense that we consider the A-GRMC simulations to be free of parameters. We also varied some of the parameters ($\tau_P$, the pausing time when two motors collide, $\tau_B$, the mean life time of the motors bound to chromatin, and $p_{on}$ and $p_{off}$) in order to assess their effect on $P(s)$ as a function of the genomic distance, $s$. The results are shown in Fig. S4

### 2 Calculation of the number of steps per second

In an earlier study (Takaki et al. 2021), we calculated the step-size distribution, $P(L|R)$, for loop-extrusion by assuming that a condensin of length $R$ captures a loop with contour length $L$. The distribution, $P(L|R)$ is given by,

$$P(L|R) = C\frac{4\pi N\{L\}(R/L)^2}{L(1-(R/L)^2)^{9/2}}\exp\Big(-\frac{3t\{L\}}{4(1-(R/L)^2)}\Big), \tag{3}$$

where $t\{L\} = 3L/2l_p$ with $l_p = 50\,nm$ is the persistence length of the DNA, and $N\{L\} = \frac{4\alpha^{3/2}e^{\alpha}}{\pi^{3/2}(4+12\alpha^{-1}+15\alpha^{-2})}$ with $\alpha\{L\} = 3t/4$. The normalization constant $C$ is independent of $L$. The distribution $P(L|R)$ does not have a well-defined mean and variance. Moreover, the Cumulative Density Function (CDF) of $P(L|R)$ cannot be easily calculated, which makes it difficult to sample $P(L|R)$ in simulations. Therefore, we numerically discretized $P(L|R)$ from $0 \leq L \leq 100,000$ nm. We sampled the discretized distribution by linearly interpolating between adjacent points in the CDF.

By using $P(L|R)$, we calculated the number of steps per second required to achieve a loop extrusion (LE) rate of $1,500bp/s$, which is the measured value in single molecule experiments at small external loads. In the experiment (Ganji et al. 2018) the LE rate = $1,500bp/s$ corresponds to 0.3 relative DNA extension. In their paper, the relative extension is defined as the length between the tethers at either end of the DNA divided by the length of unextruded DNA. From Fig. 3 in Ganji et al. 2018, the length between the tethers is $\approx 4\mu$m. Thus, for 0.3 relative extension, the length of the DNA that is not extruded is $\frac{4}{0.3} = 13.33\mu$m. Given the length of the $\lambda-$DNA in the experiment is $20\mu m$, the length of the loop is $(20 - 13.33)\mu m \approx 6.7\mu m$.

We found it takes $\approx 9.63$ steps using Eq. 3, to extrude a loop of length $6.7\mu m$, implying each step extrudes a loop with length $6.7/9.63 \approx 0.695\mu$m. Then, using an LE rate of $\approx 1500$ bp/s, and assuming that the average distance between two base-pairs is 0.34 nm, we obtain the stepping rate of condensin as 1500 x 0.34 x $10^{-3}/(0.695\mu\text{m}) \approx 0.73 \approx 1$ step/second.

The step size $L$ is chosen from the distribution Eq. 3, where $R$, the size of condensin, which we take to be $40\,nm$ (Takaki et al. 2021). We assume that when $P_{step} = 1$, both condensin I and II take one step per second to match the measured 1,500 bp/s LE velocity (Ganji et al. 2018). In most of the paper, we set $P_{Step} = 1$ and only modulate it to reproduce the experimental $P(s)$ for times, $t = 7.5, 10, 15, 30$ minutes, as the DT40 cell cycle progresses.

Following an experimental study (Kim et al. 2020), we assume that when there is a collision between different condensin sites on the chromatin (either mobile or fixed), the loop-extrusion is paused for $\tau_P \approx 7\,s$. We also varied $\tau_P$ to ascertain the impact on $P(s)$ (see Fig. S4). After $\tau_P$, the mobile sites are allowed to move through the blocks. In the LE simulations, the chromosome is modeled as a linear track with 100 bp resolution, which is less than the canonical value of the DNA persistence length. On this length scale, DNA is best described as a semi-flexible polymer, making it necessary to use the step-size distribution given in Eq. 3. The time-resolution in the simulation is $1\,s$, and the LE simulations are performed for about 60 minutes.

### 3 $P(s)$ is captured using Rouse model

The agreement between the theoretical (A-GRMC) and experimental $P(s)$ is not sensitive to the exact nature of $P(L|R)$. As shown in Fig. S1, instead of using Eq. 3, we used the $P(L|R)$ derived from $P(R|L)$ where $P(R|L)$ which for a Rouse Chain with length $L$ and monomer size $b$ has the following form,

$$P(L|R) = C \cdot 4\pi R^2 \frac{3}{2\pi \cdot L \cdot b}^{3/2} \exp\Big(-\frac{3R^2}{2 \cdot L \cdot b}\Big), \tag{4}$$

where $C = \frac{b}{6R}\{1/(1 - \text{Erf}\sqrt{\frac{3R}{2b}}))\}$ (Fig. S1(A)), where $Erf(x)$ is the error function. Interestingly, the shape of the $P(s)$ curve obtained using Eq. 4 is similar to the one obtained with the one obtained using Eq. 3.

### 4 Calculations of contact map and structures using A-GRMC

**Contact map:** We first derive an expression for the contact map for a particular choice of loop anchors (the trace of the condensin motors) specified by $L$. Let $\mathbf{r}_i = (x_i, y_i, z_i)$ be the position of the $i^{th}$ locus. The GRMC energy function associated with the loop conformation is (Bryngelson and Thirumalai 1996; Shi and Thirumalai 2021),

$$\mathcal{H} = \frac{k}{2}\sum_{i=1}^{N-1}(\mathbf{r}_{i+1} - \mathbf{r}_i)^2 + \frac{k}{2}\sum_{\alpha=1}^{n(L)-1}(\mathbf{r}_{L(\alpha+1)} - \mathbf{r}_{L(\alpha)})^2, \tag{5}$$

where, $L(\alpha)$ represents the $\alpha^{th}$ loop. The harmonic nature of the GRMC energy function makes it separable, $\mathcal{H} = \mathcal{H}_x + \mathcal{H}_y + \mathcal{H}_z$, where $\mathcal{H}_x = \frac{k}{2}\sum_{i=1}^{N-1}(x_{i+1} - x_i)^2 + \frac{k}{2}\sum_{\alpha=1}^{n(L)-1}(x_{L(\alpha+1)} -$

$x_{L(\alpha)})^2$, which in matrix notation is, $\mathcal{H}_x = \frac{1}{2}\mathbf{x}^T\mathbf{A}\mathbf{x}$. The elements, $A_{ij}$, are given by,

$$\mathbf{A}_{ij} = \begin{cases} k + \theta_i k, & \text{if } i = j = 1 \text{ or } N,\ \theta_i = 1 \text{ if i}\epsilon L \text{ and } \theta_i = 0 \text{ otherwise} \\ 2k + 2\theta_i k, & \text{if } i = j \neq 1 \text{ or } N,\ \theta_i = 1 \text{ if i}\epsilon L \text{ and } \theta_i = 0 \text{ otherwise} \\ -k, & \text{if } |i - j| = 1 \\ -k, & \text{if } i = L(\alpha - 1) \text{ and } j = L(\alpha), \text{ or } i = L(\alpha + 1) \text{ and } j = L(\alpha) \\ 0, & \text{otherwise} \end{cases} \tag{6}$$

In the first two terms of the above equation, $\theta_i$ is 1 when $i$ is a loop-base, and is 0 otherwise.

The distribution function is given by,

$$\Psi(\mathbf{r}_N) = \Psi(\mathbf{x}_N)\Psi(\mathbf{y}_N)\Psi(\mathbf{z}_N) \propto exp\left[-\frac{1}{k_BT}\left(\mathcal{H}_x + \mathcal{H}_y + \mathcal{H}_z\right)\right]. \tag{7}$$

The property of the multivariate Gaussian distributions allows us to write,

$$\Psi(\mathbf{x}_N) \propto exp\left[-\frac{1}{2k_BT}\mathbf{x}^T\mathbf{A}\mathbf{x}\right]. \tag{8}$$

The matrix $\mathbf{A}$ cannot be inverted because it is not translationally invariant, which is easily resolved by fixing the first monomer at the origin with the spring constant $k$. This modifies $\mathbf{A}$, and we can define the matrix $\widetilde{\mathbf{A}}_{ij}$ such that,

$$\widetilde{\mathbf{A}}_{ij} = \begin{cases} \mathbf{A}_{11} - k \\ \mathbf{A}_{ij}, & \text{For all other } i \text{ or } j \end{cases} \tag{9}$$

The covariance matrix is the inverse, $\Sigma = \widetilde{\mathbf{A}}^{-1}$. Thus, the equilibrium probability distribution of the x-coordinates is a normal distribution, $\widetilde{\Psi}(\mathbf{x}_N) \sim \mathcal{N}(0, \Sigma)$. The distribution of the distance along the x-axis between each pair of monomers may be written as,

$$P(x_i - x_j = x) \sim \mathcal{N}(0, \sigma_{ij}^2), \tag{10}$$

where $\sigma_{ij}^2 = \Sigma_{ii} + \Sigma_{jj} - 2\Sigma_{ij}$. Our earlier study (Shi and Thirumalai 2019) showed that the distance distribution between the $i^{th}$ and the $j^{th}$ monomer is given by,

$$P(|\mathbf{r}_{ij}| = r) = \sqrt{\frac{2}{\pi}}\frac{1}{\sigma_{ij}} e^{-r^2/2\sigma_{ij}^2}\frac{r^2}{\sigma_{ij}^2}. \tag{11}$$

The contact probability between $i$ and $j$ loci is obtained by integrating Eq. 11 till a cutoff distance, $r_c$, which yields,

$$P_{ij} = \int_0^{r_c} dr \sqrt{\frac{2}{\pi}}\frac{1}{\sigma_{ij}} e^{-r^2/2\sigma_{ij}^2}\frac{r^2}{\sigma_{ij}^2} \tag{12}$$

$$= \text{Erf}\left(\frac{r_c}{\sqrt{2}\sigma_{ij}}\right) - \sqrt{\frac{2}{\pi}} e^{-r_c^2/2\sigma_{ij}^2}\frac{r_c}{\sigma_{ij}}. \tag{13}$$

We used $r_c = b$. By repeating this calculation for all the $i$ and $j$ pairs gives us the contact probabilities for each pair of loci for a given set, $L$ of loop-bases.

**Practical implementation:** The procedure for the calculation of the loop size and the coordinates of the monomers in the loop are best explained using a single loop between the locus $i$ and $i+k$ on DNA in time $T$. The total number of steps taken is $N_{step} = \frac{T}{\Delta t}$ where $\Delta t = 1s$. At each time step a loop of length, $l_j$, which is sampled using Eq.3 with $R = 40nm$, is extruded. The loop length (in units of 10 kbps) is $k = \sum_j l_j$ where $j$ goes from 1 to $N_{step}$.

In order to compute the coordinates of all the loci between $i$ and $i+k$, we first calculated the connectivity-matrix $\mathbf{A}$ given in Eq. 6. The quadratic nature of the energy function associated with the A-GRMC enables us to compute the normal modes of the system. Each coordinate $(i, (i+1), \cdots (i+k))$ is a linear combination of the normal modes. We write $\mathbf{A}$ as $\mathbf{A} = \mathbf{U}^T \cdot \mathbf{D} \cdot \mathbf{U}$, where $\mathbf{D}$ is a diagonal matrix, and $\mathbf{U}$ is the eigenvector of $\mathbf{A}$. Furthermore, we can write $\mathbf{A} = \mathbf{U}^T \cdot \mathbf{D}^{1/2} \cdot \mathbf{I} \cdot \mathbf{D}^{1/2} \cdot \mathbf{U}$, where $\mathbf{I}$ is an identity matrix. Eq. 8 can now be rewritten as

$$\Psi(\mathbf{x}_N) \propto exp\left[-\frac{1}{2k_BT}\boldsymbol{\lambda}^T\mathbf{I}\boldsymbol{\lambda}\right], \tag{14}$$

where $\boldsymbol{\lambda} = \left(\mathbf{D}^{1/2} \cdot \mathbf{U} \cdot \mathbf{x}\right)$. Since $\mathbf{I}$ is the identity matrix Eq. 14 can be separated into $N$ independent $\mathcal{N}(0,1)$ distributions. These are the distributions of the normal mode vectors. We sample each normal mode vector separately from $\mathcal{N}(0,1)$ and use $\mathbf{x} = \left(\mathbf{D}^{1/2} \cdot \mathbf{U}\right)^{-1} \cdot \boldsymbol{\lambda}$ to generate the coordinates of the loci in the loop connecting $i$ and $i+k$. The same process is repeated for the $\mathbf{y}$ and $\mathbf{z}$ coordinates.

**Calculation of $p(s)$:** A contact map is constructed by computing $P_{ij}$ for every pair of $i, j$. The final average CM is computed by averaging over 500 different realizations of loops. From the average CM the probability of contact between two loci separated by a genomic length $s$, $P(s)$ was computed using,

$$P(s) = \frac{1}{n-s}\sum_{i=1}^{n-s}\mathbf{C}_{i,i+s}, \tag{15}$$

where $\mathbf{C}_{i,i+s}$ is the contact frequency between the $i^{th}$ and $(i+s)^{th}$ loci, and the genomic distance $s$ is measured in units of 10 kbps.

To compare between experimental and calculated $P(s)$ versus $s$, we computed the mean-squared error ($\sigma^2$) between the two $P(s)$ curves in the log scale. This is given by,

$$\sigma^2 = \frac{1}{s_{max}}\sum_{s=0}^{s_{max}}\left[\log\left[P(s)_{exp}\right)\right] - \log\left[P(s)_{GRMC}\right]\right]^2. \tag{16}$$

A small value of $\sigma^2$ implies good agreement between the two $P(s)$ curves.

#### 6 Assessing the Handedness of Mitotic Structures

In an effort to understand the helical nature of the mitotic chromosomes, we first computed $C(s)$ for synthetically generated structures. We created helices with 100 Mbps of mitotic chromsomes at 100 Kbps resolution (1,000 loci). The period of the helix ($p$) is the contour length along the helix needed to complete one turn. This corresponds to the first peak in the $C(s)$ curve (see Fig. 7 in the main text). For the synthetic helices, we used a period of 1 Mbps.

In order to elucidate the nature of helix perversion, we introduced a randomness parameter ($r$) that controls the probability with which the helix changes the handedness. In Fig. S5(A) we show the structures of the helices for different $r$ values. An ideal helix with a definite handedness through out the length of the mitotic chromosome corresponds to $r = 0$. The $C(s)$ for an ideal helix is perfectly periodic (Fig. S5(B)), which is not found in the $C(s)$ calculated using the 3D HIPPS-generated structures (see Fig. 7(B) in the main text). Moreover, $C(s)$ for a structures with Periodic Helix Perversion (PHP), is also periodic (see the purple curve in Fig. S5(B)), and inconsistent with the results in Fig. 7(B) in the main text. In contrast, for $r \neq 0$ (in particular for $r = 0.04$) the $C(s)$ obtained from the synthetic structures resembles the one calculated for condensin I depleted DT40 cells at $t = 30$min (see Fig. S5(B) in the main text), providing the first hint that helix perversion could occur randomly. The qualitative agreement between $C(s)$ calculated using the synthetic and HIPPS-generated structures for $r \neq 0$ suggests that mitotic chromosomes could also have alternating handedness, which was noted recently (Chu et al. 2020).

To further quantify the handedness of chromosomes, we used the local helical parameters ($\chi$) introduced by Moradi et al. 2009. In Fig. S6(A), we show the geometric construction for calculating $\chi$. The $\chi$ parameter depends on the period of the helix. For a helix with period $p$, $\vec{AB}$ and $\vec{CD}$ are the vectors between the $i^{th}$ and $(i + p/2)^{th}$ centers, and $(i + p/4)^{th}$ and $(i + 3p/4)^{th}$ centers, respectively. The vector $\vec{EF}$ connects the midpoints of $\vec{AB}$ and $\vec{CD}$. Using these vectors, $\chi$ is defined as the dot product of the unit vector along $\vec{EF}$ with the unit vector along $\vec{AB} \times \vec{CD}$. We computed $\chi$ along the whole structure for different values of $i$. For an ideal right handed helix, the distribution of $\chi$ is peaked at 1. Conversely for an ideal left handed helix $\vec{AB} \times \vec{CD}$ will always point away from $\vec{EF}$, and the distribution of $\chi$ is peaked at $-1$.

We computed $\chi$ for the structures with $r = 0.01$ and $r = 0.04$ (Fig. S6(C-D)). The distribution of $\chi$ for $r = 0.01$ has peaks at $-1$ and 1 in addition to 0. This implies that there are both left and right handed helices. At $r = 0.04$, which produces $C(s)$ that resembles the one shown in Fig. 7A in the main text using 3D structures, the peaks at $-1$ and 1 are lost. The distribution peaks at $\chi = 0$. At $r = 0.04$ the $C(s)$ graphs shows evidence of periodicity, but the helix is neither left or right handed but is a mixture of the two. This would correspond to helix perversion with the handedness switching randomly across the structure.

Finally, we calculated the distribution of $\chi$ using the $1,000$ structures calculated from HIPPS in condensin II chromosomes at $t = 30$ min (Fig. S6(E)). For these calculations, we used the period $p = 9.9$ Mbps, corresponding to the first peak in the $C(s)$ curve (Fig. 7(A) in the main text). The HIPPS calculations suggest that the entire ensemble of chromosomes could consist of a mixture of left and right handed helices, which is consistent with the conclusions reached previously (Zhang and Wolynes 2016). When we plot the distribution of $\chi$ for a single HIPPS structure in condensin II chromosomes at $t = 30$ min (Fig. S6(F)), we find that there is a modest bias towards $\chi < 0$. This particular structure has hints of being slightly left handed. Thus, mitotic chromosomes in general could have alternating handedness in a single structure, with possible bias towards helices with either handedness.

(A)

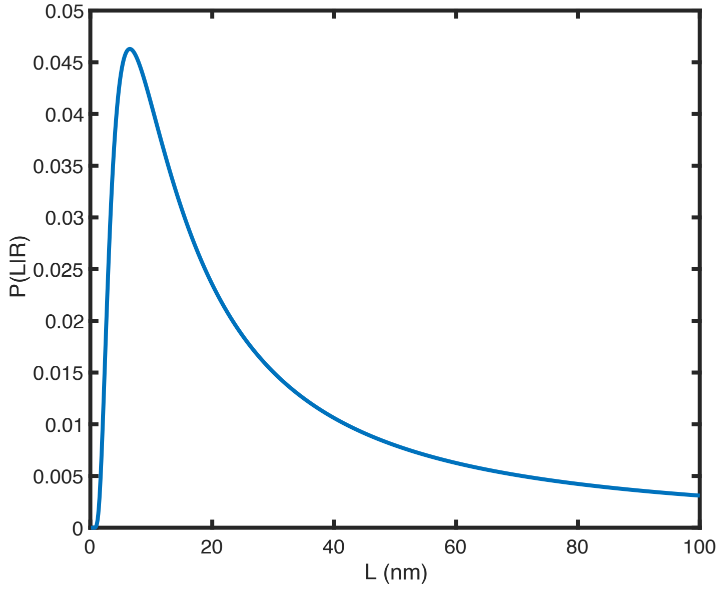


(B)

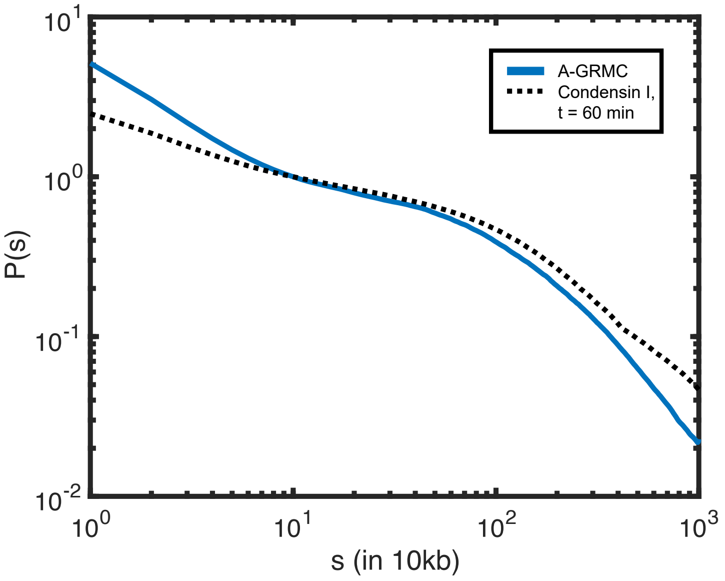


Figure S1: Calculation of $P(s)$ as a function of $s$ using Eq. 4. DNA is modelled as a Gaussian polymer with $b = 240\,nm$. The size of the loop extruded by condensin is $R = 40\,nm$. (A) Plot $P(L|R)$ using Eq. 4 shows that the distribution has a peak at $\approx 15\,nm$, which is considerably less than the persistence length of DNA. Thus, DNA is treated as a semi-flexible polymer when considering loop-extrusion by the motors. (B) Plot of $P(s)$ calculated using the step-size distribution displayed in (B). The calculated $P(s)$ (solid line) agrees well with the experimental data (Gibcus et al. 2018) (dashed line.)

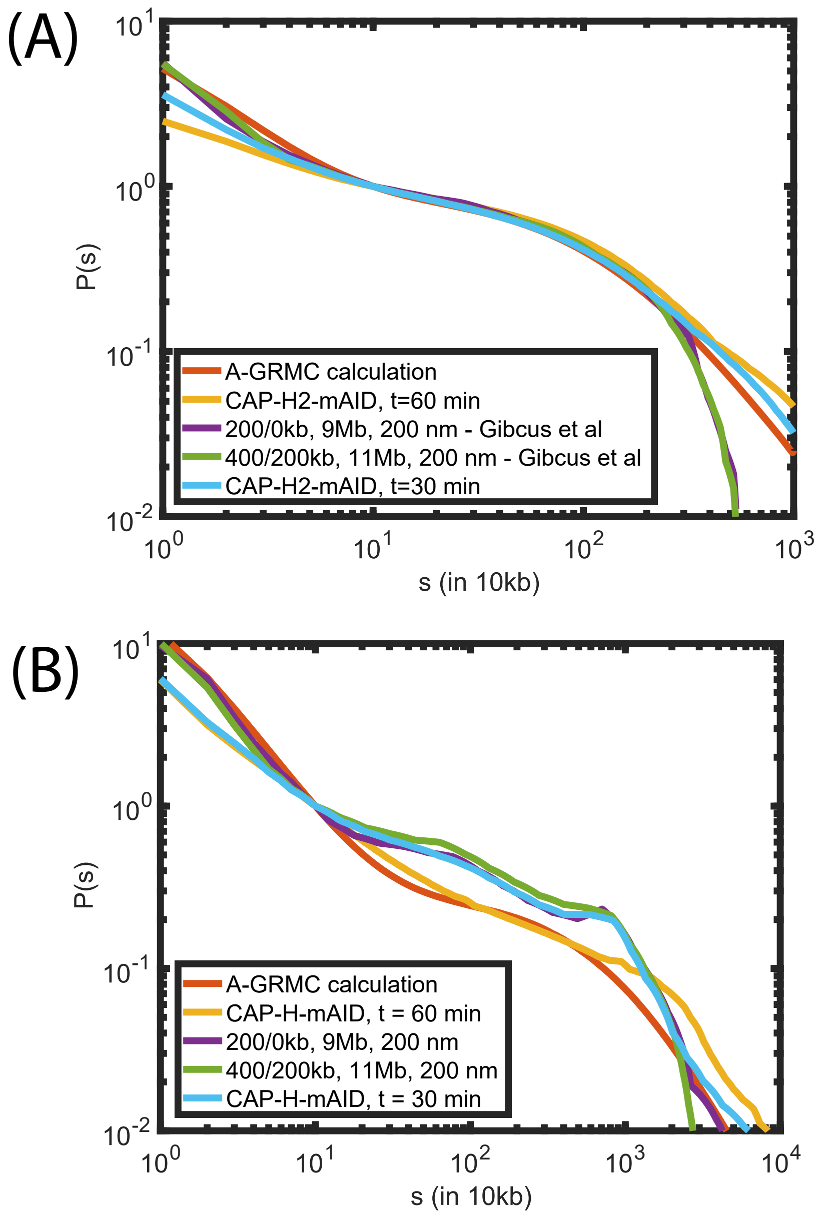


Figure S2: (A) The top panel compares $P(s)$ curves upon condensin II depletion. The A-GRMC calculations are in orange, and the experimental result at 60 minutes for the DT40 cells is in yellow. For purposes of comparison, the $P(s)$ reported by Gibcus et al. 2018 are also plotted as violet and green traces. The $P(s)$ were calculated using two sets of parameters that are specified in the figure. The light-blue is the experimental $P(s)$ at 30 minutes. (B) The bottom panel provides a similar comparison as (A) when condensin I is depleted.

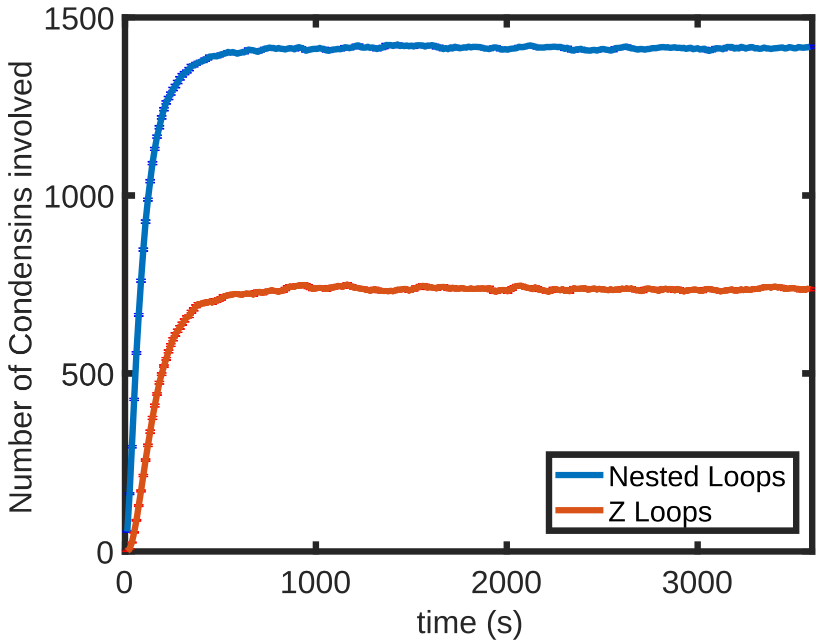


Figure S3: Total number of condensins (I and II) as a function of time in the generation of Z-loops and nested loops. A schematic of a nested loop is shown in Fig. 2C in the main text.

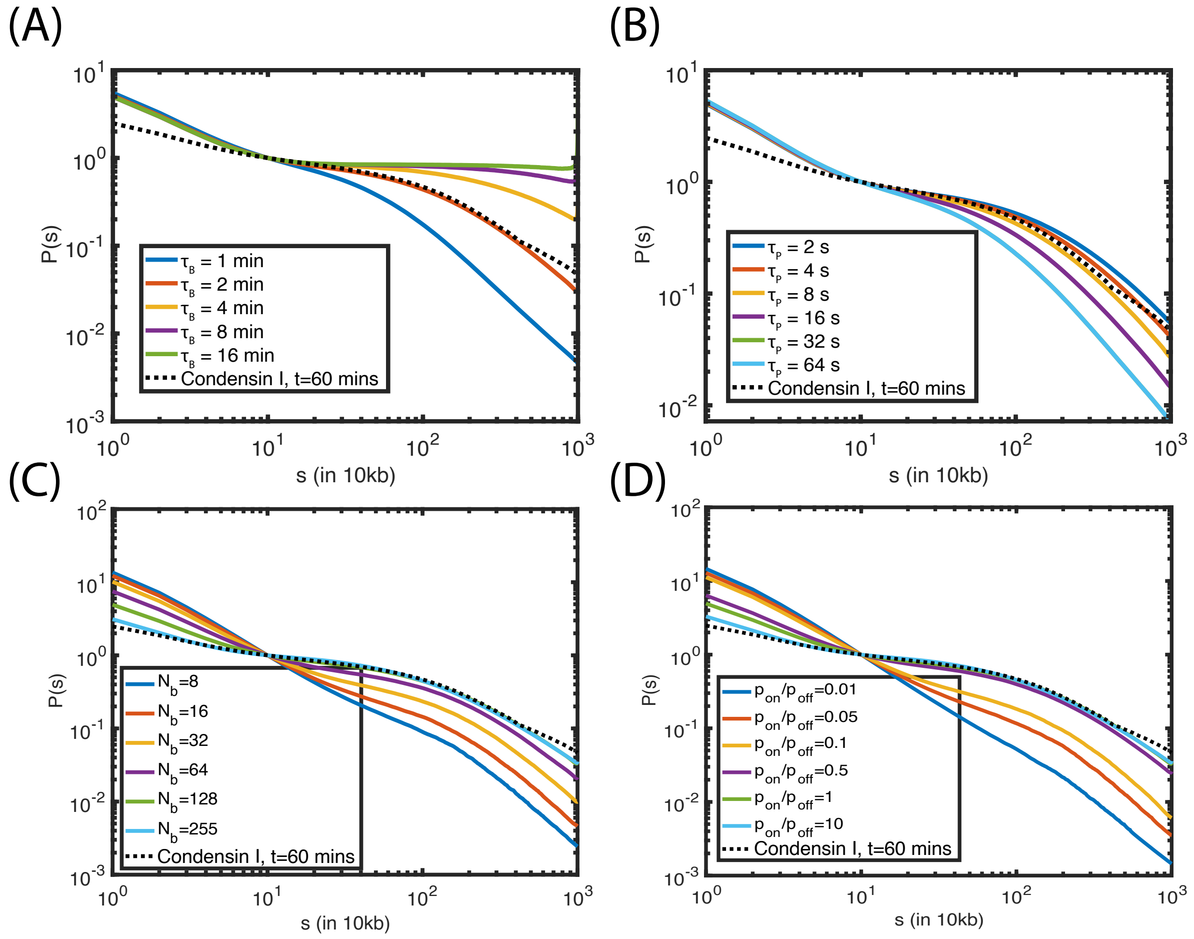


Figure S4: **Changes in $P(s)$ upon variation of condensin parameters in the A-GRMC model (see Table 1 in the main text)**. The results are for condensin I with the total chromatin length being 10 Mbps. (A) Increasing the $\tau_B$, the duration of the time the motor is bound to DNA, increases $P(s)$ at high $s > 10^5$ bps. (B) Increasing the pausing time $\tau_P$ decreases $P(s)$ because the average loop-length decreases. (C) Increasing the number of bound condensins also increase the $P(s)$ at high s, until it saturates at the maximum number of condensins available. (D) Increasing $p_{on}/p_{off}$ enhances long range contacts while depleting the shorter range contacts. This is reflected in an increase in $P(s)$ at high $s$.

**Table S1: Total number of CAP-H proteins present in the cell extracted from, Graph A in Fig. S2(Walther et al. 2018) using WebPlotDigitizer(Rohatgi 2017)**

| Time (in min) | Number of CAP-H |
|---|---|
| 1.3 | 386791 |
| 4.1 | 410808 |
| 7.1 | 416294 |
| 10.1 | 408979 |
| 33.7 | 398190 |
| Average | 403395 |

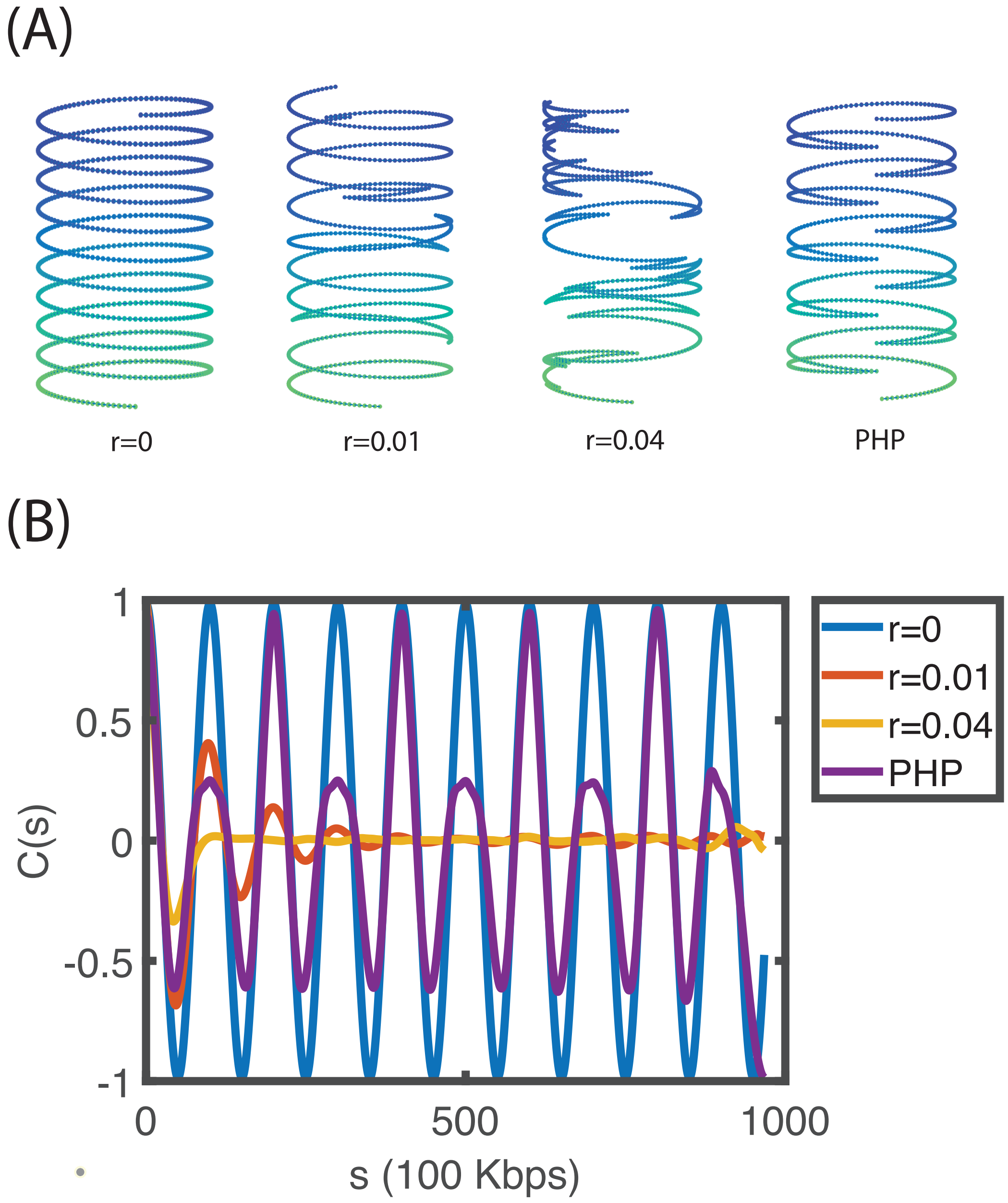


Figure S5: **Angular correlation for synthetic structures.** (A) Conformations of helices for different values of $r$. In PHP the handedness changes in a symmetric manner. (B) Calculated $C(s)$ as function of $s$ for the synthetic structures in (A). As $r$ increases there is a loss in coherence in $C(s)$ as $s$ increases. The $C(s)$ for the PHP has two different periods. The $C(s)$ structure at $r = 0.01$ resemble the $C(s)$ for condensin 2 chromosomes at $t = 30$ min, whereas, $r = 0.04$ resembles the $C(s)$ for HeLa cells. The colors are explained in the box on the right.

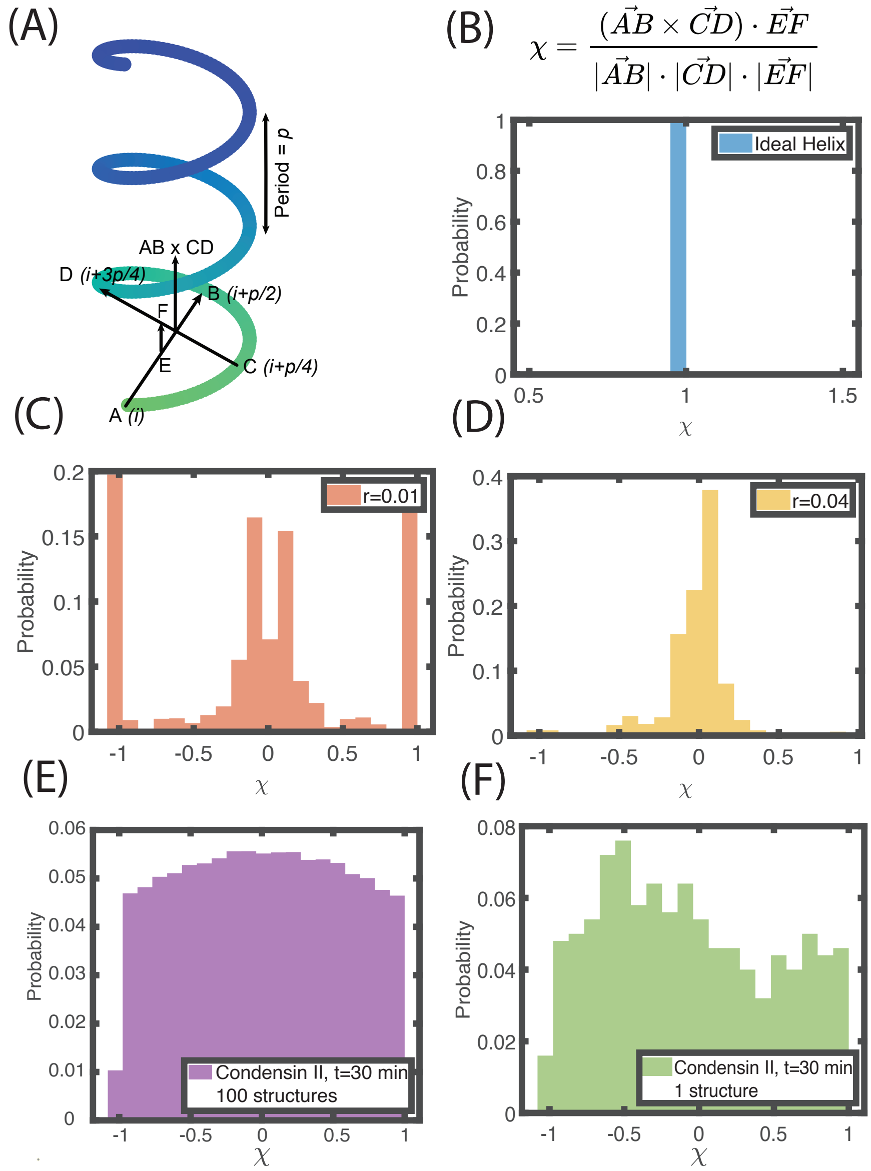


$$\chi = \frac{(\vec{AB} \times \vec{CD}) \cdot \vec{EF}}{|\vec{AB}| \cdot |\vec{CD}| \cdot |\vec{EF}|}$$

Figure S6: **Distribution of local handedness.** (A) Geometric construction used to calculate the handedness parameter ($\chi$). For a helix with period $p$, $\vec{AB}$ and $\vec{CD}$ are vectors between the $i^{th}$ and $(i + p/2)^{th}$ loci, and $(i + p/4)^{th}$ and $(i + 3p/4)^{th}$ loci respectively. The vector $\vec{EF}$ connects the midpoints of $\vec{AB}$ and $\vec{CD}$. (B) Mathematical definition of $\chi$. For an ideal right handed helix, the distribution of $\chi$ is peaked at 1. (C) The distribution of $\chi$ for a helix with perversion for $r = 0.01$ has peaks at $-1$ and 1 but also at 0. (D) At $r = 0.04$ the peaks at $-1$ and 1 are lost whereas the peak at 0 is prominent. (E) The distribution of $\chi$ averaged over $1,000$ structures calculated using HIPPS in condensin II chromosomes at $t = 30$ min. For these calculations, we used the period $p = 9.9$ Mbps. The structures exhibit both left and right handed helices. (F) The $\chi$ distribution for HIPPS structure in condensin II chromosomes at $t = 30$ min shows that it is possible to have a slight bias for either right or left handed helix within a single structure.

**Table S2: Total number of CAP-H2 proteins present in the cell extracted from, Graph A in Fig. S2(Walther et al. 2018) using WebPlotDigitizer(Rohatgi 2017)**

| Time (in min) | Number of CAP-H2 |
|---|---|
| 4.9 | 54219 |
| 36.8 | 54219 |
| Average | 54219 |

**Table S3: Total number of CAP-H proteins bound to DNA extracted from, Graph C in Fig. 1(Walther et al. 2018) using WebPlotDigitizer(Rohatgi 2017)**

| Time (in min) | Number of CAP-H |
|---|---|
| 0.9 | 67317 |
| 2.6 | 89416 |
| 3.6 | 116050 |
| 5.2 | 133699 |
| 7.0 | 139919 |
| 8.9 | 143180 |
| 20.1 | 142996 |
| 22.0 | 143088 |
| 23.8 | 144011 |
| 25.7 | 145026 |
| 27.5 | 146318 |
| 29.4 | 151487 |
| 30.0 | 154863 |
| 31.3 | 164500 |
| 33.1 | 188680 |
| 35.0 | 190433 |
| 36.5 | 167216 |
| 37.5 | 151117 |
| 38.7 | 158685 |
| 40.5 | 143273 |
| Average | 151066 |

**Table S4: Total number of CAP-H2 proteins bound to DNA extracted from, Graph C in Fig. 1(Walther et al. 2018) using WebPlotDigitizer(Rohatgi 2017)**

| Time (in min) | Number of CAP-H2 |
|---|---|
| 1.5 | 38087 |
| 2.4 | 36623 |
| 3.3 | 35372 |
| 4.2 | 34629 |
| 5.1 | 33675 |
| 6.0 | 33059 |
| 29.3 | 32699 |
| 30.2 | 32975 |
| 31.1 | 33038 |
| 32.0 | 33038 |
| 32.9 | 32805 |
| 33.8 | 32593 |
| 34.7 | 32063 |
| 35.6 | 32020 |
| 36.5 | 32020 |
| 37.2 | 30366 |
| Average | 33050 |
